# An *in silico* evaluation of anthraquinone derivatives as potential inhibitors of DNA gyrase B of *Mycobacterium tuberculosis*

**DOI:** 10.1101/2022.10.01.510436

**Authors:** Juliana C. Amorim, Andrea E. Cabrera Bermeo, Viviana E. Vásquez Urgilés, Maritza R. Martínez León, Juan M. Carpio Arévalo

## Abstract

The World Health Organization reported that tuberculosis remains on the list of the top ten threats to public health worldwide. Among the main causes is the limited effectiveness of treatments due to the emergence of a diverse spectrum of resistant strains of *Mycobacterium tuberculosis*, the causative agent of the disease. One of the main drug targets studied to combat *M. tuberculosis* is DNA gyrase, the only enzyme responsible for regulating DNA topology in this specie and considered essential in all bacteria. In this context, the present work tested the ability of 2,824 anthraquinones retrieved from the PubChem database to act as competitive inhibitors through interaction with ATP-binding pocket of DNA gyrase B of *M. tuberculosis*. Virtual screening results based on molecular docking identified 7122772 (N-(2-hydroxyethyl)-9,10-dioxoanthracene-2-sulfonamide) as the best-scored ligand. From this anthraquinone, a new derivative was designed harbouring an aminotriazole moiety, which exhibited higher binding energy calculated by molecular docking scoring and free energy calculation from molecular dynamics simulations. In addition, in these last analyses, this ligand showed to be stable in complex with the enzyme and further predictions indicated an acceptable pharmacokinetic profile. Taken together, the presented results show a new anthraquinone synthetically accessible and with promising potential to inhibit the GyrB of *M. tuberculosis*.

## Introduction

Tuberculosis is a mostly respiratory disease caused by *Mycobacterium tuberculosis*.^1^ The disease currently affects about 10 million people worldwide, with an average of 1.3 million deaths per year.^2^ Even with the introduction of the BCG vaccine more than a century ago, the global rate of infection with *M. tuberculosis* is one in three people.^3^ In addition, the bacteria have the ability to escape from the cells of the immune system and remain dormant in old lesions which result in ineffective and prolonged treatment.^3, 4^ The treatment currently standardized for World Health Organization as first-line against drug-susceptible *M. tuberculosis* infection is for six months and is effective in approximately 85% of cases. ^2, 4^ It initially consists of the administration of ethambutol, isoniazid, pyrazinamide and rifampicin for two months, followed by four months using isoniazid and rifampicin.^3, 4^ Among the main causes that prevent a successful treatment are the incorrect prescription of drugs or the lack of them in the health units, and also the low adherence to treatment by patients. ^2^ As a consequence, multidrug-resistant strains of *M. tuberculosis* emerge and adapt, which become more efficient in the infective process and the evolution of tuberculosis.^5^

The increasing accessibility of next-generation sequencing technologies has allowed considerable progress in the understanding of the dynamics between different *M. tuberculosis* strains and the host during infection.^6^ Furthermore, it became possible to continuously extract information about mutations in the essential drug targets and adapt or develop new treatment options. ^5, 7^ Among the targets considered essential for bacterial survival validated for antituberculosis drug discovery is DNA gyrase.^8^ This enzyme is representative of the type II topoisomerases in bacteria and is responsible for catalyzing the introduction of negative DNA supercoils in an ATP-dependent manner.^9^ The main activities of this enzyme are related to events of replication, transcription, and also recombination.^10^ However, in *M. tuberculosis*, besides these functions, this enzyme is the unique responsible for decatenation, which in other bacteria is carried out by DNA topoisomerase IV. ^11^ DNA gyrase is composed of two subunits called GyrA and GyrB, which form an A2B2 heterotetramer when active. While GyrA is responsible for breaking and reunion of the DNA strands, GyrB promotes the hydrolysis of ATP to make energy available for DNA supercoiling. ^9,10^ Among the main inhibitors described for GyrA are quinolones (e.g., ciprofloxacin), antibiotics responsible interfering with the bacterial DNA-gyrase-DNA complex.^12^ On the other hand, some of the main inhibitors described for GyrB have affinity for the ATP-binding pocket, acting competitively with ATP and thus preventing its hydrolysis.^13^ Among the main classes of inhibitors of this target are drugs such as aminocoumarins,^14^ but there are also a wide variety of promising molecules tested *in silico*, *in vitro*, and *in vivo* assays such as aminopyrazinamides, ^15^ bithiazoles,^16^ indazoles,^17^ pyrrolamides,^18^ and also pyrazoltiazoles. ^19^ However, a large proportion of the described inhibitors for GyrB have not been successful in clinical trials. From this limitation, it becomes necessary to expand the search for new inhibitors for this target. Compounds that show promising potential for a large number of therapeutic applications are phytochemicals, ^20-23^ which are structurally diverse and complex groups of molecules. Among this class, the anthraquinones are scaffolds described as potential inhibitors of nucleotide-binding proteins.^24^ These molecules represent the largest group of naturally occurring quinones and are structurally defined by the presence of a central benzoquinone between two benzene rings, which may be functionalized by various types of chemical groups.^25^ Based on the above, the present work evaluated through *in silico* approaches the affinity of 2,824 anthraquinones for the ATP-binding pocket of *M. tuberculosis* GyrB (*Mt*GyrB) to identify the most suitable chemical substituents to develop potential inhibitors based on the anthraquinone backbone.

## Methods

### Target and ligands preparation

For the present work was used the *Mt*GyrB model named 3ZKBL (RCSB PDB crystal 3ZKB, resolution: 2.9 Å, free R-value: 0.240, working R-value: 0.182, observed R-value: 0.184,^26^), which has the 216-239 loop region computationally rebuilt as described in a previous publication.^27^ 3ZKBL was prepared using the Dock Prep module of the UCSF Chimera-1.16,^28^, then processed by the SPORES 1.3 tool using default parameters and saved in MOL2 format.

The chemical structures of the 2, 824 anthraquinones were obtained from the PubChem database (https://pubchem.ncbi.nlm.nih.gov/) and downloaded in SDF format in March 2022. This database was generated from the substructure of the anthraquinone backbone and applying the following filters: rotatable bonds: maximum 10, hydrogen bond donors: maximum 5, hydrogen bond acceptors: maximum 10 and in the data source category: chemical vendors. Finally, these molecules were filtered to include anthraquinones with molecular below 350 g/mol to restrict the search to aglycones and exclude dimeric structures and heterosides. The co-crystallized ligand, ANP, (33113 ([[[[(2R,3S,4R,5R)-5-(6-aminopurin-9-yl)-3,4-dihydroxyoxolan-2-yl]methoxy-hydroxyphosphoryl]oxy-hydroxyphosphoryl]amino]phosphonic acid) was used as control. Subsequently, hydrogen atoms were assigned to the structures at pH 7.4 for all the ligands, the 3D structure was generated, followed by minimization steps of these structures with the MMFF94 force field,^29^ using the steepest descent geometry optimization with 500 steps, followed by conjugate gradient algorithm with default parameters and transformed into MOL2 format, all using Open Babel-3.1.1 software.

### Molecular docking-based virtual screening analyses

The molecular docking-based virtual screening analyses were performed using PLANTS-1.2 software.^30^ A 20 Å radius centering the coordinates of the ATP-binding pocket of 3ZKBL was set to X= −26.86, Y= −27.11, and Z= 17.77. The search speed was set to one and the scoring function was selected as ChemPLP. The clustering RMSD was set to 2.0 Å and all docking scores were calculated by the default scoring function.

### Pharmacokinetic and synthetic accessibility predictions

To obtain information on the pharmacokinetic properties of the most promising molecules and also of the reference compounds, predictions were made using the Swiss-ADME server.^31^ The properties evaluated were gastrointestinal absorption, permeability across the blood-brain barrier, potential transport by P-gp (P-glycoprotein) and the likelihood of interactions associated with cytochromes related to drug metabolism.

Theoretical prediction of the synthetic accessibility of the promising compounds was performed using the Ambit-SA software (http://ambit.sourceforge.net/reactor.html). The algorithm of the software uses four different descriptors to represent different structural and topological features: the molecular complexity, the steric chemical complexity, the complexity due to the presence of fused and also bridged systems. The results are presented as scores ranging from 0 to 100, where the value 100 represents the most easily synthesizable molecule.

For both described analyses, SMILES codes of the molecules were generated and used as input files.

### Target fishing predictions

Given that off-target effects of drugs can provoke adverse effects,^32^ and knowing that anthraquinones interact with several human proteins,^33^ it was decided to perform a target fishing analysis to evaluate comparatively the ability of the most promising molecules and of the reference compounds to interact with human targets. The results are presented as scores ranging from 0 to 1, where the value 1 represents the major probability of being targets of each molecule. For the prediction of potential targets of these compounds the SMILES codes of the molecules were generated and used as input files to the Swiss Target Prediction web server.^34^

### Molecular dynamic simulations

Molecular dynamics simulations (MDS) were performed using the GROMACS-2021.1 software,^35^ to better understand the interactions and stability of 3ZKBL and their complexes with selected ligands. All simulations were performed using the CHARMM 36 all-atom force field.^36^ The transferable water model of intermolecular potential 3P was used for solvation by selecting a periodically corrected cubic box using a minimum distance of 1 nm. Later, the system was neutralized by adding Na^+^ and Cl^ˉ^ ions, to eliminate the initial steric shocks 100,000 energy minimization steps were conducted using the steepest descent algorithm. The system was equilibrated during 500 and 5,000 ps at 310 K and 1 bar pressure in the NVT and NPT arrays, respectively. The runs were conducted during 100 ns and the coordinates were saved every 10 ps. The Particle Mesh Ewald algorithm,^37^ was used to analyze the long-range electrostatic interactions and the LINCS algorithm,^38^ was used to regulate the covalent bonds. The Leap-frog algorithm and the Berendsen coupling,^39^ were used to control the pressure and temperature.

### Binding free energy calculation

To complement the molecular docking and MDS analyses, were performed free energy calculations among 3ZKBL and the selected ligands on a single trajectory based on the MMPBSA method,^40^ using the gmx-mmpbsa 1.5.2 software.^41^ To perform the calculations was extracted the MDS results from the last 50 ns of the runs using 2,500 snapshots for each simulation. The parameters that were used to calculate the free energies were inp=1, istrng=0.15, and indi=2, the other parameters were kept according to the software recommendations.

### Visualizations of molecular docking and molecular dynamics results

The structures of the novel anthraquinones were designed using the Marvin JS web server (https://marvinjs-demo.chemaxon.com/latest/). 2D interaction diagrams of protein-ligand complexes were generated with the PoseView web server.^42^ The 3D diagrams were visualized using the software Discovery Studio Visualizer-2021 (https://discover.3ds.com/). MDS analyses were visualized with GROMACS scripts in conjunction with Python scripts using the NumPy, Pandas, Matplotlib, Seaborn and Pytraj libraries. The RMSD, RMSF and Rg analyses were generated from the alpha-carbon of the protein in the presence or absence of the ligands, while the hydrogen bonds were generated from the protein-ligand complexes.

## Results and Discussion

### Virtual screening of anthraquinones

The search for *Mt*GyrB inhibitors has been a collaborative effort of academia and industry over many decades.^13^ Research groups have isolated a lot of natural compounds and also conducted the synthesis of a wide variety of inhibitors of this target.^43, 44^ In parallel, there have also been considerable advances in the availability of *Mt*GyrB crystal structures,^26^ more complete and with better resolution, which have also allowed advances *in silico* studies to the discovery of potential inhibitors.^45, 46^

In the present work, 2,824 anthraquinones, identified by its PubChem ID, were ranked based on their best scores from molecular docking-based virtual screening (Supplementary Table 1). Among the ten best scores (BS), presented in descending order of binding affinity are BS-1 (7122772, N-(2-hydroxyethyl)-9,10-dioxoanthracene-2-sulfonamide), BS-2 (85992490, (9, 10-dioxoanthracen-2-yl) methanesulfonate), BS-3 (3794639, 1-[(5-nitropyridin-2-yl)amino]anthracene-9,10-dione), BS-4 (133109345, (1,4-dihydroxy-9,10-dioxoanthracen-2-yl) hydrogen sulfate), BS-5 (94940647, 1-(9, 10-dioxoanthracen-2-yl)triazole-4-carboxylic acid), BS-6 (108793505, N-(9,10-dioxoanthracen-1-yl)-6-oxo-1H-pyridazine-3-carboxamide), BS-7 (7122774, N,N-dimethyl-9,10-dioxoanthracen-2-sulfonamide), BS-8 (625367, 6-chloro-9,10-dioxoanthracene-2-carboxylic acid), BS-9 (100282, 1,4-bis(2-aminoethylamino)anthracene-9,10-dione), BS-10 (123132836, (E)-3-[5-(9,10-dioxoanthracen-1-yl)furan-2-yl]prop-2-enoic acid). As can be seen in Table 1, of these molecules, six derivatives have substitutions at position 2, of which four have sulfur-oxygen containing groups, such as sulfonamide (BS-1 and BS-7), sulfonate (BS-2) and sulfate (BS-4). Another three anthraquinones have substitution at position 1 (BS-3, BS-6 and BS-10) and one have groups in position 1 as well as in position 4 (BS-9).

**Table 1.**
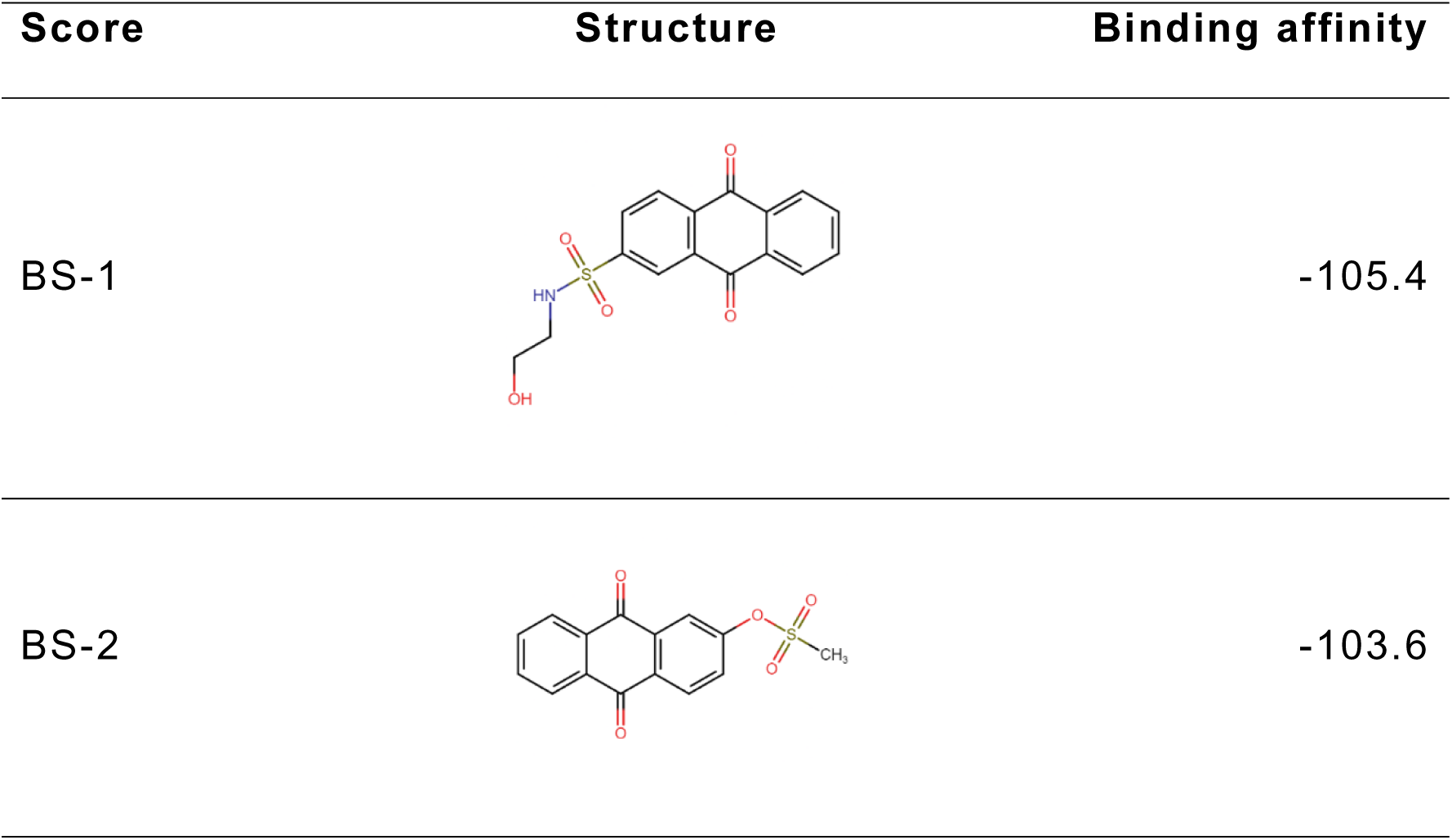
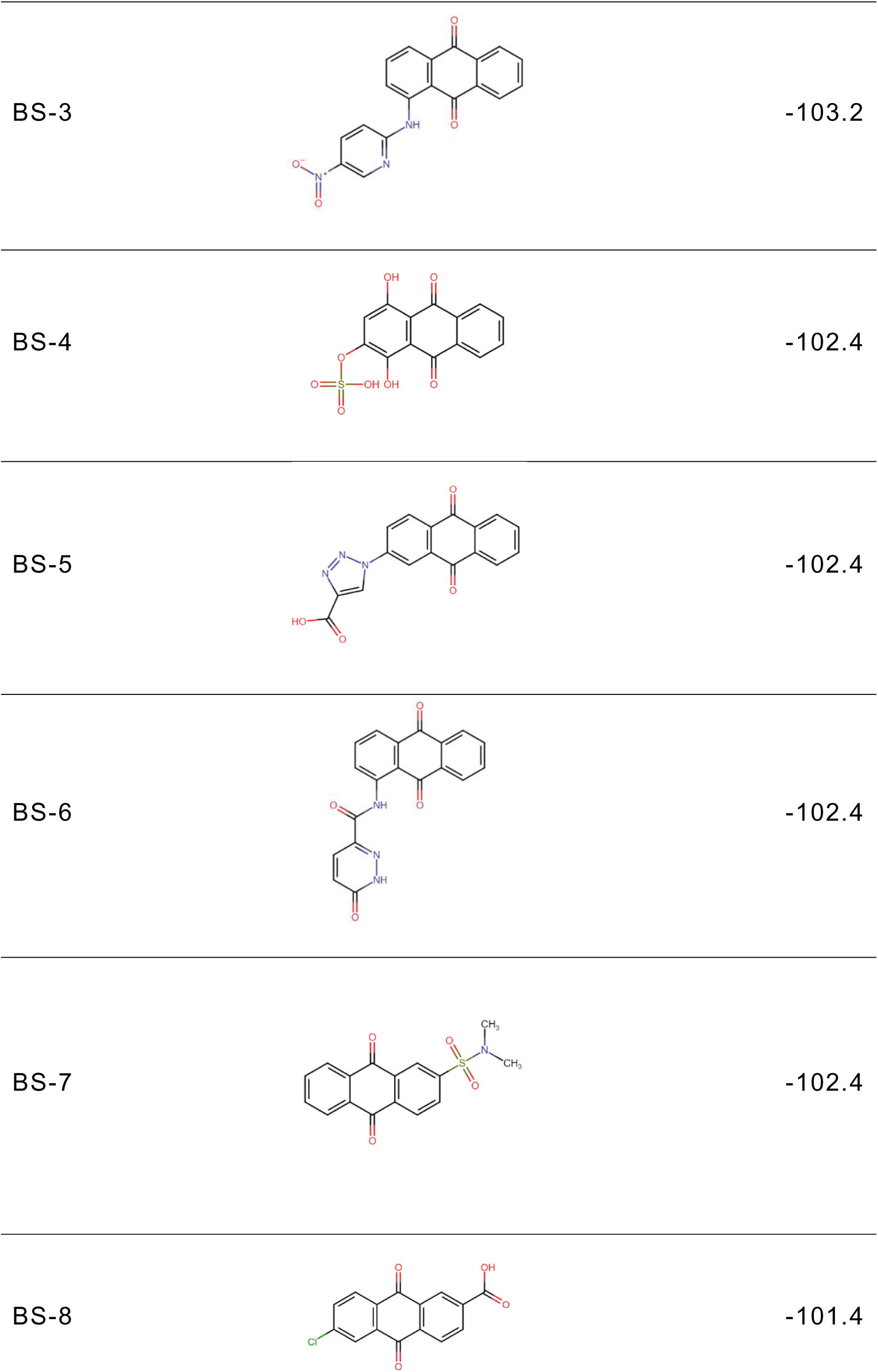
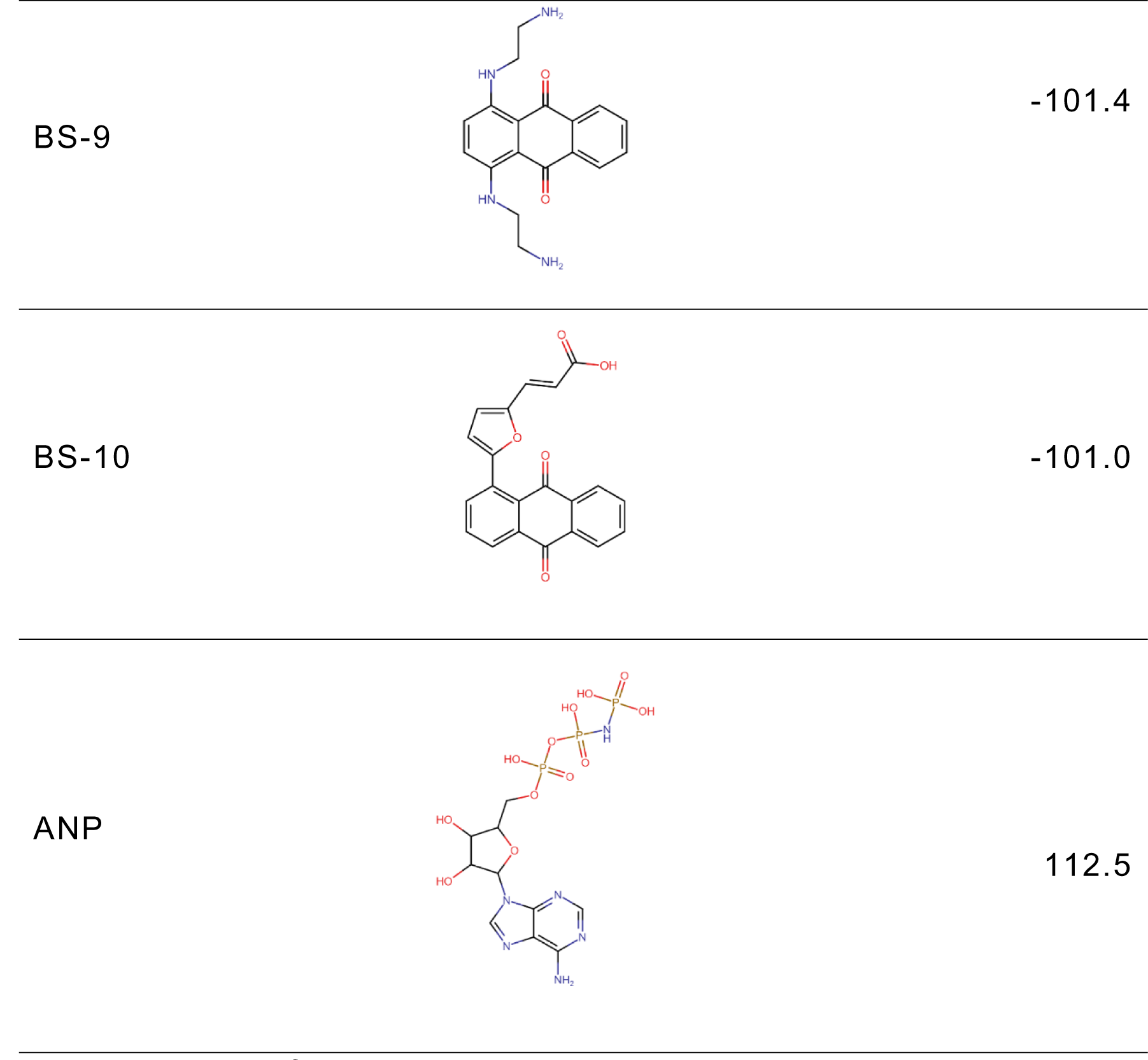
Structure and binding affinities of the best-scored anthraquinones. BS-1 (7122772), BS-2 (85992490), BS-3 (3794639), BS-4 (133109345) and, BS-5 (94940647), BS-6 (108793505), BS-7 (7122774), BS-8 (625367), BS-9 (100282), BS-10 (123132836) and ANP (33113). These analyses were performed with PLANTS software.

Importantly, all of these substituents include hydrogen bond donor and acceptor groups. Although the binding affinity of BS-1 (−105.4) and the aforementioned anthraquinones is slightly lower than that of ANP (−112.5), their polar substituents are oriented towards the interior of the catalytic site, analogous to that of the phosphate groups of the co-crystallized ligand (Table 1 and Fig.1).

**Fig. 1.**
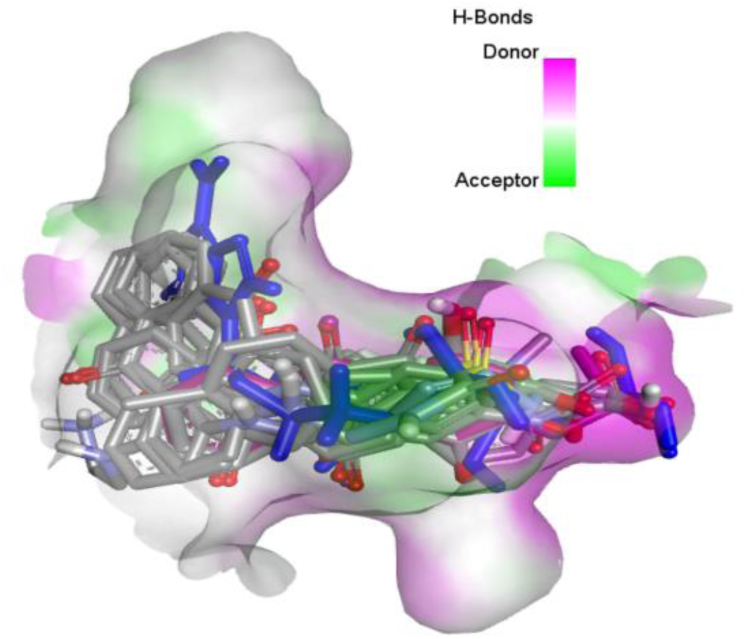
Superposition of the best poses of the 10 best-scored anthraquinones compared to crystallographic pose of ANP in the catalytic site of MtGyr B. BS-1 (7122772), BS-2 (85992490), BS-3 (3794639), BS-4 (133109345), BS-5 (94940647), BS-6 (108793505), BS-7 (7122774), BS-8 (625367), BS-9 (100282) and BS-10 (123132836). ANP is highlighted in dark blue. 3D hydrogen bond surface representation was performed with Discovery Studio Visualizer.

Considering the higher binding affinity and the representativeness of sulfur-oxygen-containing groups among the aforementioned anthraquinones, the sulfonamide derivative, BS-1, was selected for further analysis.

### Comparative analyses of the ANP and BS-1 poses

As shown in the Fig.2A, BS-1 formed several hydrogen bonds with the amino acids Asn52, Gly107, Val123, Gly124 and Val125 of 3ZKBL. With the exception of interaction with Gly107, all these hydrogen bonds are also established among the ANP and 3ZKBL (Fig.3A). The Fig.2B shows that the sulfonamide linked to the hydroxyethyl substituent of BS-1 projects deeply into the ATP-binding pocket like that of the phosphate groups of the ANP (Fig.3B). However, the number of interactions formed by BS-1 in the catalytic site of 3ZKBL is lower compared to the interactions of ANP with this enzyme. In addition, while ANP forms a hydrogen bond between the amino group of its adenine and the carboxylate group of Asp79 (Fig.3A), which is a key residue to the bacterial GyrB function,^47, 48^ BS-1 shows no interaction with this amino acid (Fig.3B).

**Fig. 2.**
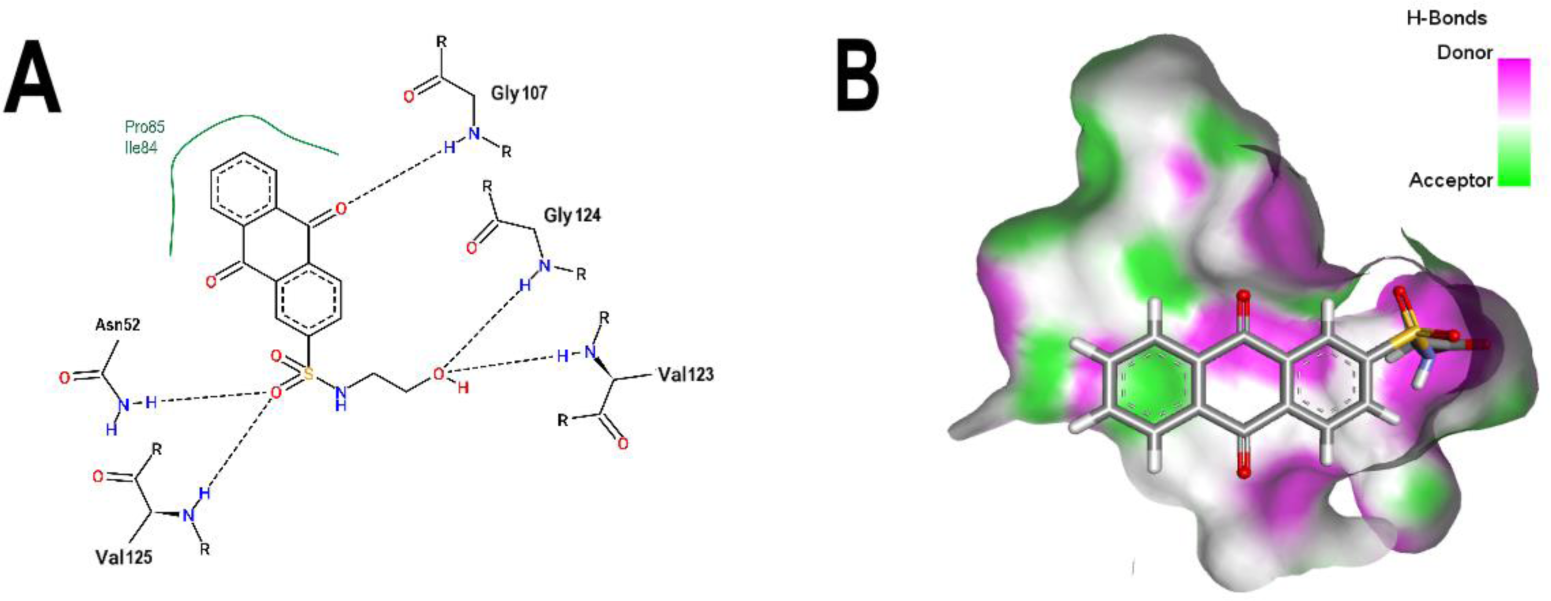
Analysis of the best pose of BS-1. A) 2D interaction diagram of the 3ZKBL-BS-1 complex. B) 3D representation of hydrogen bond surface of 3ZKBL-BS-1 complex. 2D interaction diagram was performed with PoseView. 3D hydrogen bond surface representation was performed with Discovery Studio Visualizer.

**Fig. 3.**
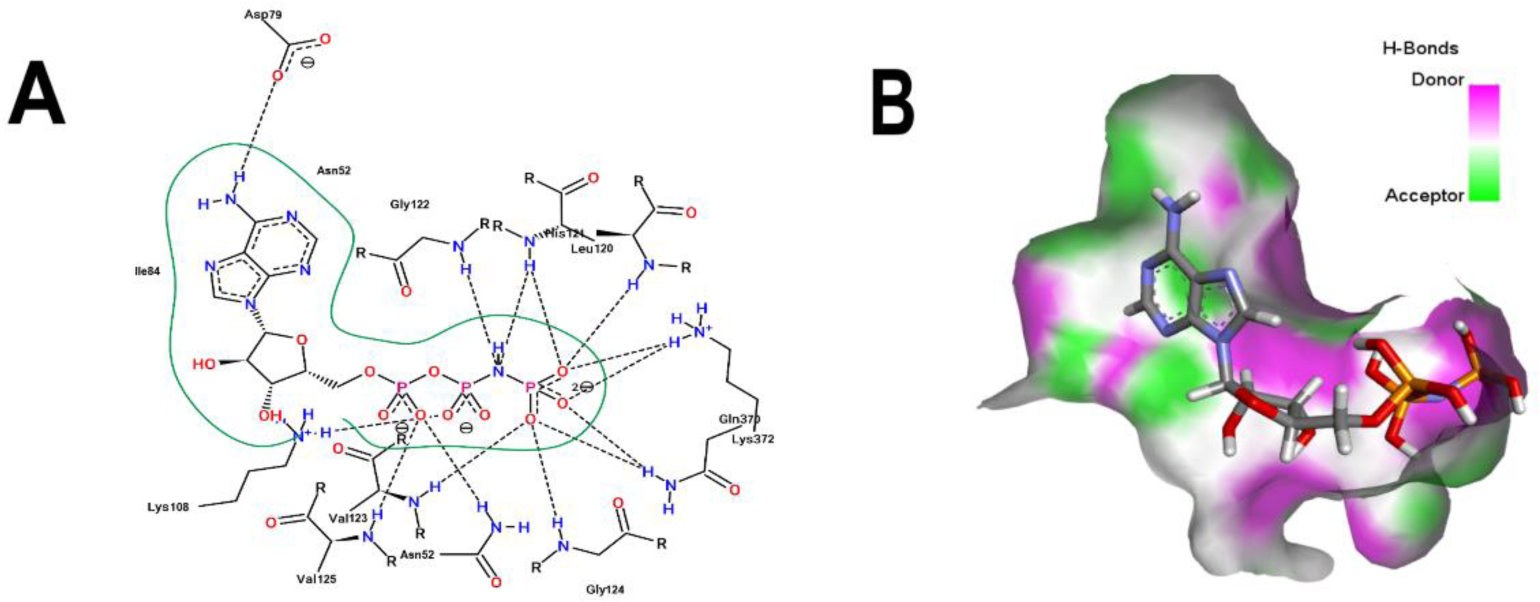
Analysis of the crystallographic pose of ANP. A) 2D interaction diagram of 3ZKBL-ANP complex. B) 3D representation of hydrogen bond surface representation 3ZKBL-ANP complex. 2D interaction diagram was performed with PoseView. 3D hydrogen bond surface representation was performed with Discovery Studio Visualizer.

From these observations and to resemble the above-mentioned interaction between BS-1 and 3ZKBL, an aminotriazole group located on carbon eight of its anthraquinone backbone was introduced, generating a new molecule (hereinafter called ATD). This moiety is part of the structure of 1-(3-amino-1H-1,2,4-triazol-5-yl)anthracene-9,10-dione (PubChem CID: 153843066), although there is no previous information of the potential biological activities of this molecule. The rationale for this modification was to fill the region in the active site occupied by the adenine of ANP co-crystallized, which is flanked by hydrogen bond acceptor groups including the carboxylate of Asp79 (Fig.2B and 3B). To evaluate the synthetic accessibility of this new derivative was used the Ambit-SA software, which show a score of 60.41 indicating that ATD is a synthesizable anthraquinone.

Fig.4 shows that the amino group on the triazole ring attains the objective of forming the hydrogen bond with Asp79, which provoke only a small shift in the RMSD values of its backbone compared to that of the parent molecule (0.342 Å). Indeed, except for Gly107, ATD form the same interactions than BS-1. In addition, this new aminotriazole derivative also exhibited an increased binding affinity to reach a value of −107.5 in PLANTS score.

**Fig. 4.**
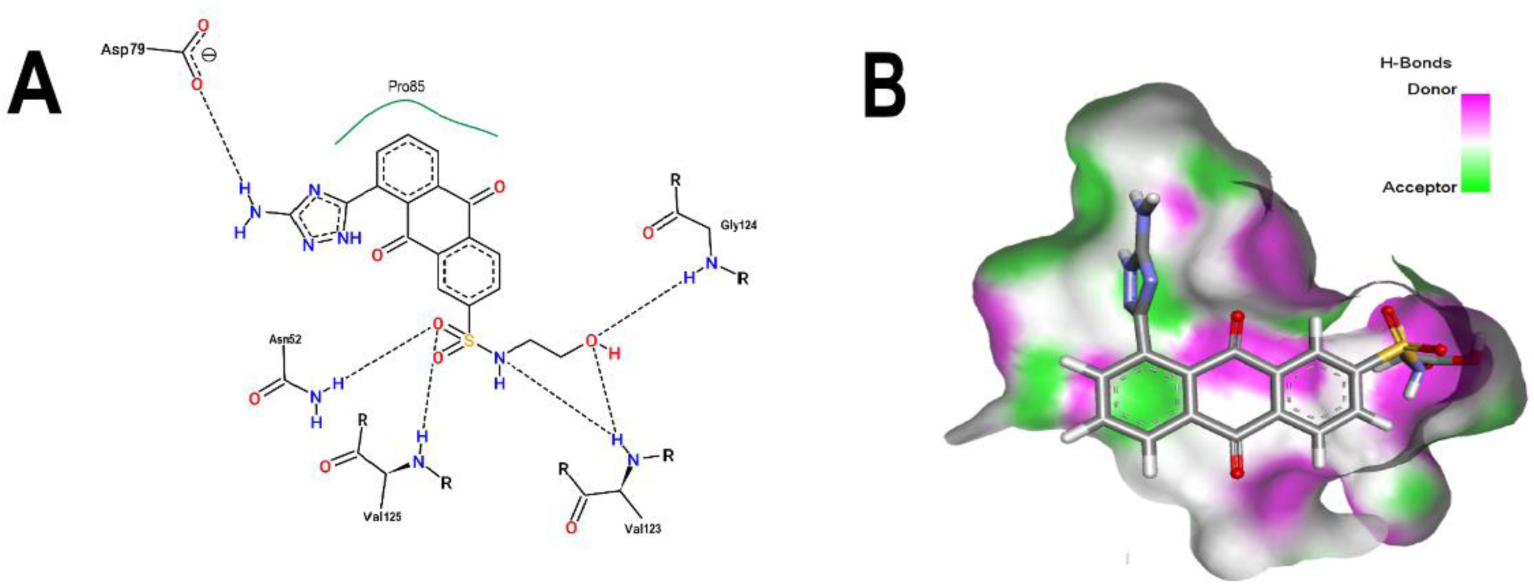
Analysis of the best pose of ATD. A) 2D interaction diagram of the 3ZKBL-ATD complex. B) 3D representation of hydrogen bond surface representation of 3ZKBL-ATD. 2D interaction diagram was performed with PoseView. 3D hydrogen bond surface representation was performed with Discovery Studio Visualizer.

### Pharmacokinetic analyses

To gain hints of some of the pharmacokinetic properties of this new derivative, predictions were made using the SwissADME server. Additionally, for comparative purposes, these same predictions were performed for the anthraquinone without functional groups (hereinafter called AC) and for the parent molecule BS-1. The predictions reveal that ATD has lower gastrointestinal absorption than AC and BS-1 and is a P-gp substrate, thus its parenteral administration may be necessary for further *in vivo* experiments. In addition, unlike AC, ATD does not have properties to cross the blood– brain barrier (BBB), which could reduce the risk of adverse effects at the central nervous system level. Importantly, contrary to AC, ATD does not act as an inhibitor of cytochromes involved in the metabolism of several drugs, which would avoid potential effects arising from interactions with known substrates of these enzymes. Altogether, these analyses show that ATD exhibits a pharmacokinetic profile suitable for further *in vivo* experiments.

### Target fishing analysis

To identify possible human targets for ATD, prediction analyses were performed using the Swiss Target Prediction server and these results were compared with the same predictions performed for AC and BS-1 (Supplementary Table 2). The results in Table 3 show that, compared to the high probability of AC to interact with the phosphatase Cdc25B (0.898), which is involved in the activation of cyclin-dependent kinases at key stages of the cell cycle,^49^ neither BS-1 or ATD would show this ability. In fact, according to this prediction tool, there is no similarity of ATD with ligands of the 3,068 proteins included in its analyses, which would suggest a low probability of ATD to interact with human targets. However, further analysis will be needed to confirm these results.

**Table 2.** Pharmacokinetic analyses of AC, BS-1 and ATD. These analyses were performed with SwissADME server.

| Pharmacokinetic analyses |  |  |  |
| --- | --- | --- | --- |
|  | AC | BS-1 | ATD |
| **GI absorption** | High | High | **Low** |
| **BBB permeant** | Yes | No | **No** |
| **P-gp substrate** | No | No | **Yes** |
| **CYP1A2 inhibitor** | Yes | No | **No** |
| **CYP2C19 inhibitor** | Yes | No | **No** |
| **CYP2C9 inhibitor** | No | No | No |
| **CYP2D6 inhibitor** | No | No | No |
| **CYP3A4 inhibitor** | No | No | No |

**Table 3.** Target fishing analyses of AC, BS-1 and ATD. The results show the top five targets of AC, BS-1 and ATD ranked according to their probability of being targets of each molecule. Probability values range from 0 to 1. Target fishing analyses were performed with Swiss Target Prediction.

| **Target fishing analyses** | **Probability** |
| --- | --- |
| **Targets of AC** |  |
| Dual specificity phosphatase Cdc25B | 0.89883662 |
| Acyl coenzyme A: cholesterol Acyltransferase 1 | 0.06042459 |
| Apoptosis regulator Bcl-2 | 0.06042459 |
| Casein kinase I alpha | 0.06042459 |
| Caspase-3 | 0.06042459 |
| **Targets of BS-1** |  |
| Thymidine kinase cytoplasmatic | 0.10161385 |
| Thymidine kinase mitochondrial | 0.10161385 |
| Alkaline phosphatase | 0.10161385 |
| P-glycoprotein 1 | 0.10161385 |
| Tyrosine-protein kinase ABL | 0.10161385 |
| **Targets of ATD** |  |
| 3-phosphoinositide dependent protein kinase-1 | 0 |
| 78 kDa glucose-regulated protein | 0 |
| Adenosine A1 receptor | 0 |
| Adenosine A2a receptor | 0 |
| Adenosine A3 receptor | 0 |

### Molecular dynamic simulations analyses

The MDS was performed to evaluate the stability and interactions of the complexes formed between 3ZKBL and the selected ligands. The first analysis was performed to evaluate the stability of apo-3ZKBL by calculating its RMSD, which showed a mean value of 0. 4 nm, with fluctuations between 0.3 and 0.6 nm throughout the simulation, which could be attributed to the flexibility of the loop in the absence of ligand. Importantly, the 3ZKBL-BS-1, 3ZKBL-ANP and 3ZKBL-ATD complexes show lower RMSD values compared to the apo form throughout the run t ime. All complexes exhibited an average RMSD of approximately 0.2 nm (Fig. 5A), which reveals that these ligands form stable complexes with the enzyme.

**Fig 5.**
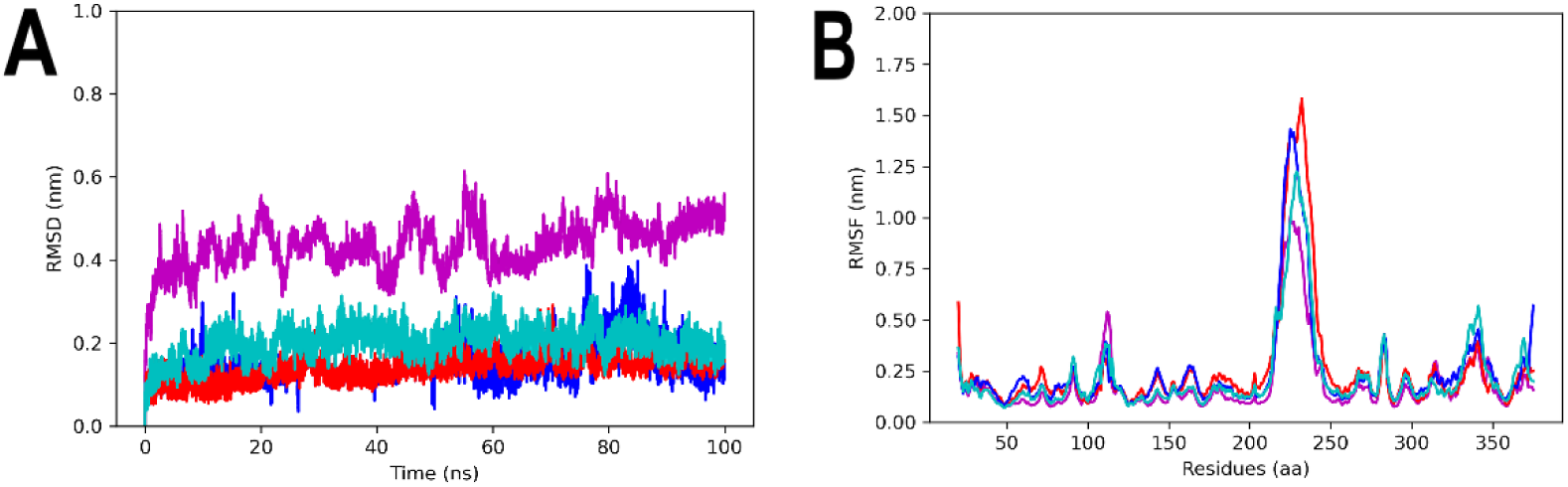
MDS analyses of apo-3ZKBL and 3ZKBL complexed with ANP, BS1, and ATD. A) RMSD values. B) RMSF values. The colours magenta, red, dark blue and light blue represent apo-3ZKBL, 3ZKBL-ANP, 3ZKBL-BS-1 and 3ZKBL-ATD respectively. The analyses were performed with Gromacs.

To identify the amino acids with higher fluctuations of apo-3ZKBL and also of its complexes were calculated their RMSF values. The results show more flexibility at residues 216 −239 (a region that corresponds to the rebuilt loop). The maximum values of RMSF corresponding to these residues were of 1.0, 1.3, 1.4, and 1.6 nm, respectively for apo-3ZKBL, 3ZKBL-ATD, 3ZKBL-BS-1 and 3ZKBL-ANP. However, it was observed that the loop containing the residues responsible for interacting with the ATP (amino acids 103-122), showed greater flexibility in apo-3ZKBL (RMSF of 0.5 nm) compared to 3ZKBL-ANP (RMSF of 0.4 nm) and to 3ZKBL-BS-1 and 3ZKBL-ATD which showed almost the same value (RMSF of 0.35 nm), Fig. 5B. These results also indicate that the ligands stabilize the enzyme, and also that the greatest fluctuations are in regions corresponding to loops, where a high flexibility in GyrB is already expected.^50^

Analyzing the Rg values of each of the systems over the simulation time, it can be seen that the complexes, as well as the apo-3ZKBL form, have very similar compaction values with average Rg values of 2.4 nm throughout the run. The maximum Rg value of apo-3ZKBL and 3ZKBL-ATD was 2.55 and 2.57 nm, respectively from the initial 5 ns of simulation. For the 3ZKBL-ANP and 3ZKBL-BS-1 complexes, the maximum Rg values were 2.47 and 2.45 nm, respectively near the 10 ns of the simulation (Fig. 6A).

**Fig. 6.**
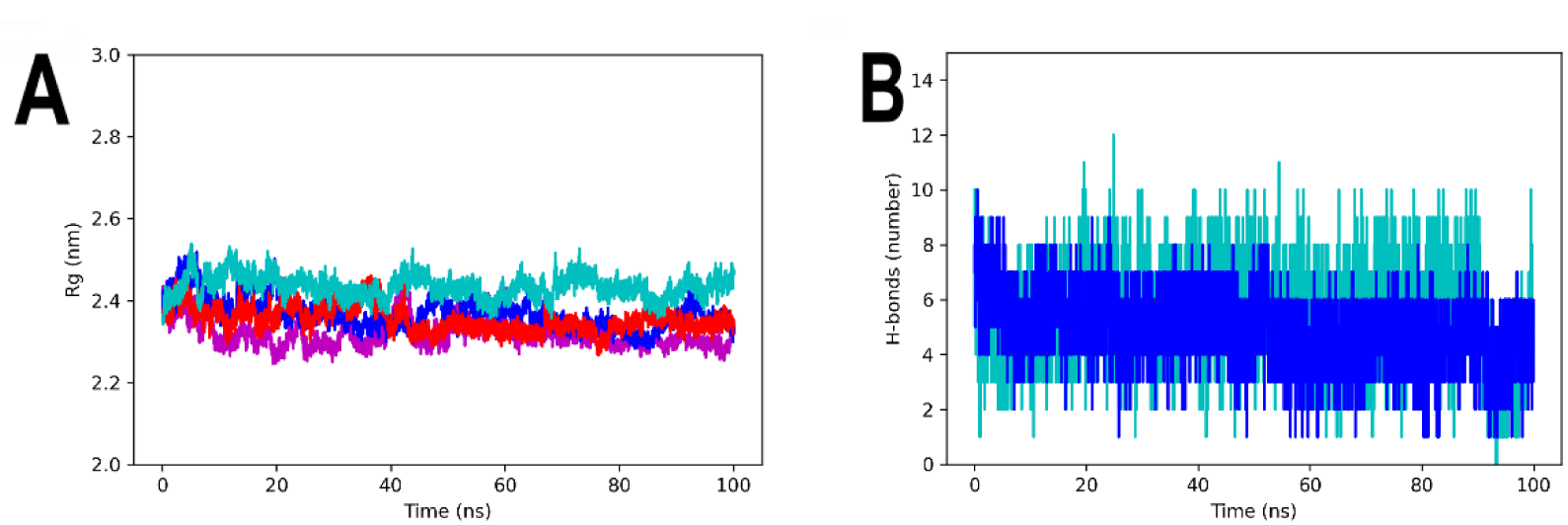
MDS analyses of apo-3ZKBL and 3ZKBL complexed to ANP, BS1, and ATD A) Rg values. B) Hydrogen bonds number. The colors magenta, red, dark blue and light blue represent apo-3ZKBL, 3ZKBL-ANP, 3ZKBL-BS-1 and 3ZKBL-ATD respectively. The analyses were performed with Gromacs.

Regarding the hydrogen bonds established among the enzyme and the ligands, it can be observed that 3ZKBL and BS-1 establish a maximum of 10 hydrogen bonds at the beginning of the run, while an average of six to eight interactions occurred intermittently during the remaining run time. While the complex formed between 3ZKBL and ATD establishes up to 12 hydrogen bonds in 25 ns, followed by 9 and 10 bonds maintained intermittently over the remaining analysis time, with the exception of the 90 to 97 ns interval where hydrogen bonds dropped to an average of three for this complex (Fig. 6B).

In addition, the formation of two or three hydrogen bonds between the Asp79 of 3ZKBL and the aminotriazole moiety of the ATD is also confirmed during most of the run (Fig. 7A), which can also be observed in the snapshot obtained in the 100 ns of the run presenting two hydrogen bonds (Fig. 7B).

**Fig. 7.**
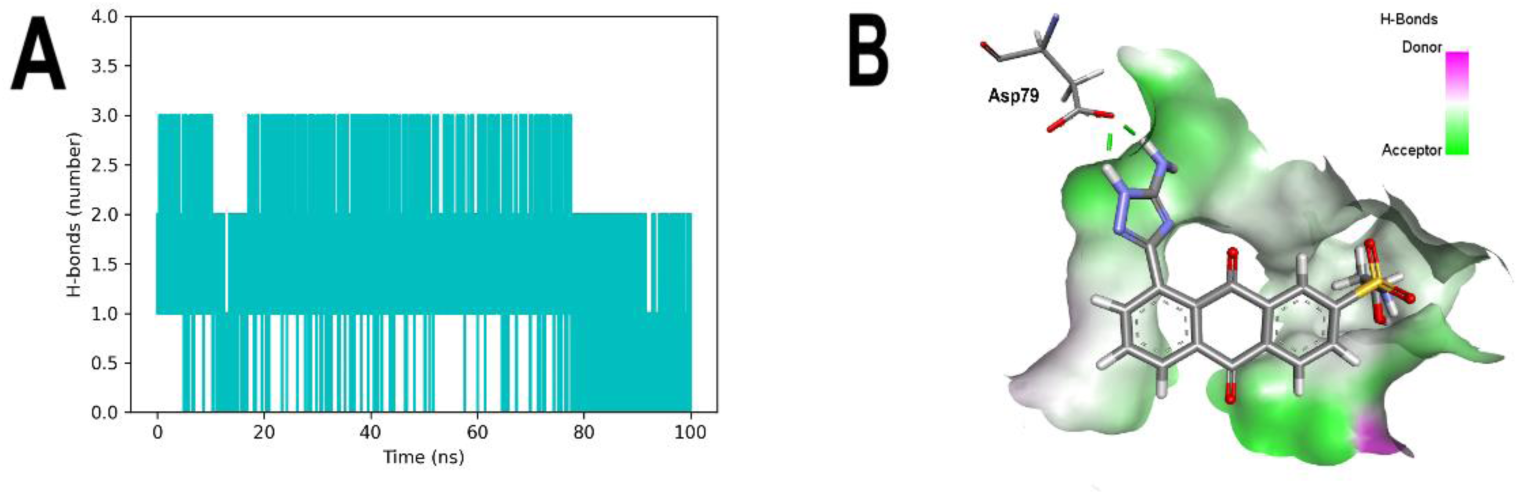
MDS analyses of 3ZKBL complexed to ATD. A) Hydrogen bonds between Asp79 and ATD in the 3ZKBL-ATD complex. B) Snapshot of 100 ns time of 3ZKBL-ATD complex highlighting the hydrogen bonds of ATD with Asp79. The hydrogen bond analysis was performed with Gromacs. 3D hydrogen bonds surface representation and hydrogen bonds detection in the snapshot at the 100 ns was performed with Discovery Studio Visualizer.

### Binding free energy calculation

The results of free energy calculations for the two complexes studied, presented in Table 4, demonstrate that there was an increment of free energy established between 3ZKBL-ATD compared to 3ZKBL-BS-1, with ΔG (Total) values of −36.48 and −17.89 kcal/mol, respectively. The decomposition of the binding energy shows an increment due to Van der Waals ΔE (Vdw) interactions, with values of −35.65 and - 48.60 kcal/mol for the complex with BS-1 and ATD, respectively. The electrostatic interactions in the complex with BS-1 present unfavorable contributions with ΔE (Ele) of 89.21 kcal/mol, while ATD showed values of −20.51 kcal/mol. It was also observed that the polar solvation energy shows favorable values in BS-1, ΔE (Polar, Solv) of - 67.43 kcal/mol and unfavorable ones in ATD with a value of 37.38kcal/mol, it was therefore presented opposite to the results of the electrostatic interactions. Overall, the results of the energy calculation corroborate the higher binding energy of ATD compared to BS-1, which is consistent with that observed in the molecular docking analyses.

**Table 4.**
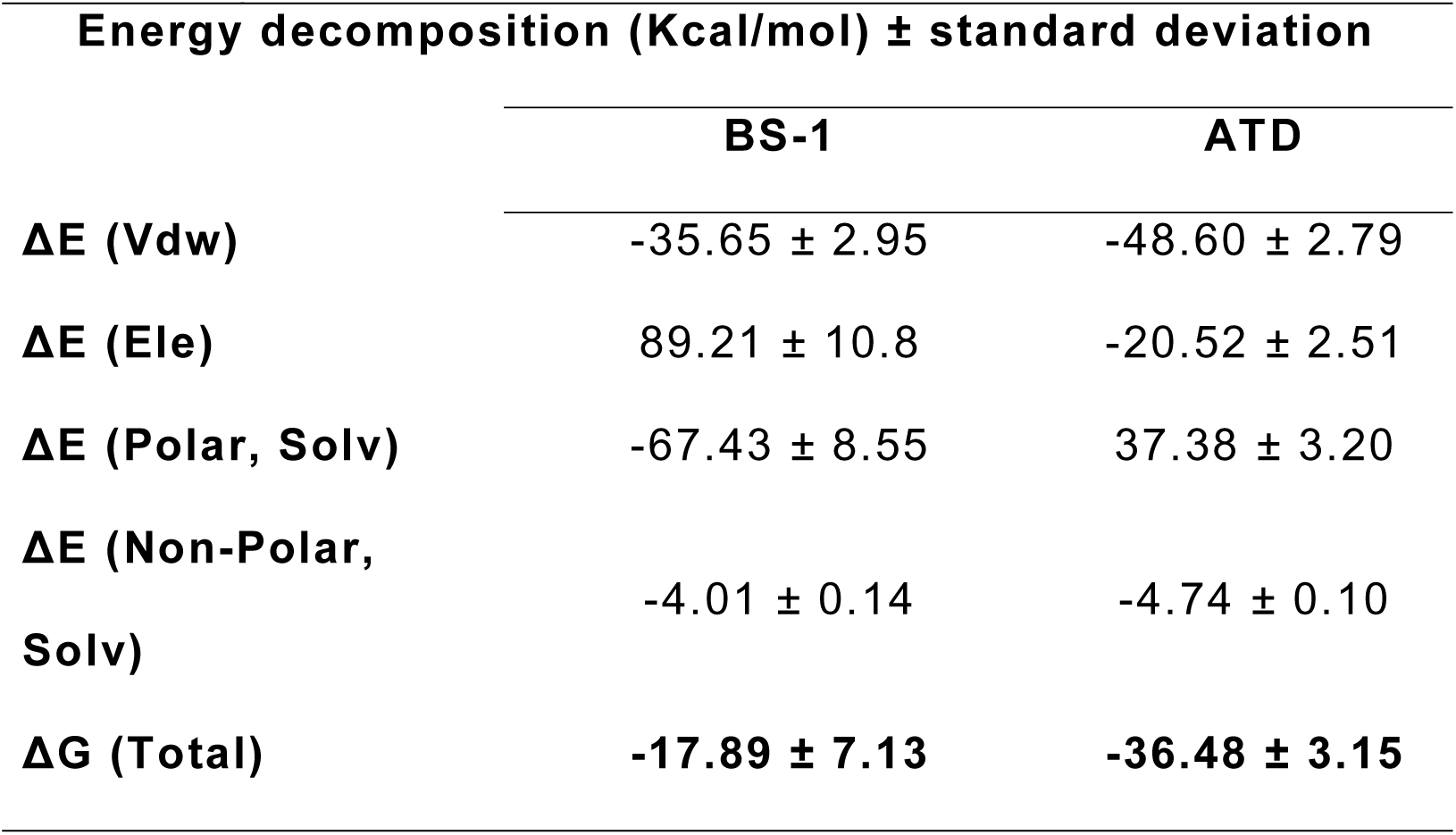
Binding free energy calculations extracted from the single trajectory of MDS analyses of the 3ZKBL-BS-1 and 3ZKBL-ATD complexes. The analyses were performed with gmx-mmpbsa software

| **Energy decomposition (Kcal/mol) ± standard deviation** |  |  |
| --- | --- | --- |
|  | **BS-1** | **ATD** |
| **$\Delta E$ (Vdw)** | -35.65 ± 2.95 | -48.60 ± 2.79 |
| **$\Delta E$ (Ele)** | 89.21 ± 10.8 | -20.52 ± 2.51 |
| **$\Delta E$ (Polar, Solv)** | -67.43 ± 8.55 | 37.38 ± 3.20 |
| **$\Delta E$ (Non-Polar, Solv)** | -4.01 ± 0.14 | -4.74 ± 0.10 |
| **$\Delta G$ (Total)** | **-17.89 ± 7.13** | **-36.48 ± 3.15** |

## Conclusion

As *M. tuberculosis* remains a threat to human health, mainly due to the limited effectiveness of existing treatments to circumvent multidrug-resistant events, the search for new molecules that can act as potential antibiotics is needed. In this work, the ability of 2,824 anthraquinones to inhibit *Mt*GyrB through association with the ATP-binding pocket was analyzed. The molecule 7122772 (N-(2-hydroxyethyl)-9,10-dioxoanthracene-2-sulfonamide, BS-1) exhibited the best binding energy for *Mt*GyrB of the entire dataset. However, the aminotriazole anthraquinone derivative (ATD), which interacts with the key Asp79 of *Mt*GyrB, outperformed BS-1 in binding energy, pharmacokinetic profile and also exhibits a lesser probability of off-target effects. In addition, in MDS analyses, ATD complexed with *Mt*GyrB show high stability and the free energy calculations exhibit better inhibitory potential than BS-1. Collectively, the results obtained in the present work reveal ATD as a new potential *Mt*GyrB inhibitor synthetically accessible based on the anthraquinone backbone.

## Author contribution

JCA, AECB, VEVU, MRML and JMCA work on methodology, validation, formal analyses, investigation and interpretation of the findings. JCA and JMCA also work on funding acquisition, conceptualization, visualization and wrote the manuscript. AECB, VEVU, MRML critically revised the manuscript. All authors approved the final version of the manuscript.

---

**Supplementary table 1.** Molecular docking-based virtual screening of 2,424 anthraquinones on the nucleotide-binding pocket of MtGyrB

|  |  |
| --- | --- |
| **Name** | **PLANTS score** |
| 7122772 | -105.422 |
| 85992490 | -103.6168 |
| 3794639 | -103.2347 |
| 133109345 | -102.451 |
| 94940647 | -102.435 |
| 108793505 | -102.432 |
| 7122774 | -102.422 |
| 625367 | -101.457 |
| 100282 | -101.439 |
| 123132836 | -101.021 |
| 71320149 | -100.408 |
| 71379713 | -100.386 |
| 71320150 | -100.031 |
| 7113016 | -99.9941 |
| 66591685 | -99.9753 |
| 7148329 | -99.7602 |
| 4385343 | -99.6674 |
| 67030 | -99.2043 |
| 82227886 | -99.0363 |
| 188986 | -98.7069 |
| 7064888 | -98.6243 |
| 9840968 | -98.6018 |
| 236903 | -98.4855 |
| 94971665 | -98.4347 |
| 16800485 | -98.0859 |
| 16807410 | -97.9634 |
| 11002466 | -97.7754 |
| 6612581 | -97.4695 |
| 112474645 | -97.4593 |
| 4263770 | -97.3654 |
| 3090908 | -97.32 |
| 235576 | -97.1633 |
| 3266379 | -97.1626 |
| 2749859 | -96.7448 |
| 3592817 | -96.7263 |
| 71364709 | -96.4904 |
| 11630782 | -96.4606 |
| 3904435 | -96.4161 |
| 22984 | -96.3937 |
| 333261 | -96.0672 |
| 2785186 | -95.9938 |
| 3085736 | -95.9495 |
| 4356619 | -95.8105 |
| 12307088 | -95.803 |
| 15700795 | -95.4401 |
| 3843618 | -95.2026 |
| 3928593 | -95.1688 |
| 2837522 | -95.1621 |
| 51062962 | -94.9935 |
| 5299796 | -94.8504 |
| 331636 | -94.8224 |
| 19926672 | -94.8197 |
| 355491 | -94.7692 |
| 3076446 | -94.6028 |
| 3633126 | -94.4224 |
| 2933577 | -94.3439 |
| 20317509 | -94.3403 |
| 5737051 | -94.2998 |
| 3090914 | -94.2754 |
| 57620624 | -94.1945 |
| 2748062 | -94.1142 |
| 433369 | -94.0003 |
| 9972777 | -93.9184 |
| 22608864 | -93.862 |
| 638297 | -93.8063 |
| 84372733 | -93.7619 |
| 12531988 | -93.7187 |
| 4342362 | -93.7009 |
| 71368256 | -93.6683 |
| 105526 | -93.5782 |
| 631257 | -93.5648 |
| 3015518 | -93.5411 |
| 10380616 | -93.5403 |
| 3465426 | -93.5207 |
| 9967100 | -93.511 |
| 3851856 | -93.4528 |
| 23680288 | -93.4449 |
| 13211699 | -93.4036 |
| 99940 | -93.3764 |
| 2749856 | -93.3362 |
| 626812 | -93.3018 |
| 14089067 | -93.2315 |
| 3076423 | -93.227 |
| 14472278 | -93.1902 |
| 3918212 | -93.1858 |
| 101945587 | -93.1205 |
| 82200641 | -93.0218 |
| 13248406 | -92.9951 |
| 8552 | -92.9711 |
| 3969677 | -92.9516 |
| 5197942 | -92.9463 |
| 24206299 | -92.8901 |
| 82316607 | -92.845 |
| 195787 | -92.8438 |
| 147677 | -92.8234 |
| 94940653 | -92.7948 |
| 46782876 | -92.7515 |
| 91885118 | -92.6784 |
| 49695398 | -92.6298 |
| 3076425 | -92.6261 |
| 374712 | -92.6019 |
| 7072925 | -92.571 |
| 16832109 | -92.567 |
| 18877474 | -92.5319 |
| 38357220 | -92.4576 |
| 54719453 | -92.4571 |
| 137163277 | -92.4461 |
| 20462886 | -92.2943 |
| 25906 | -92.2404 |
| 107849 | -92.1345 |
| 331637 | -92.1335 |
| 23521871 | -92.0943 |
| 2889110 | -92.0934 |
| 221014 | -92.0926 |
| 3085265 | -92.0649 |
| 89984 | -92.0198 |
| 71445369 | -91.9592 |
| 2871166 | -91.9396 |
| 110832078 | -91.9242 |
| 87341 | -91.8922 |
| 54678683 | -91.7958 |
| 10425624 | -91.7907 |
| 3089372 | -91.7811 |
| 3026899 | -91.7463 |
| 90158 | -91.6897 |
| 432860 | -91.6877 |
| 3966634 | -91.6682 |
| 66522 | -91.6643 |
| 7699272 | -91.6607 |
| 3090900 | -91.6514 |
| 112479887 | -91.6459 |
| 18877476 | -91.6166 |
| 625575 | -91.4954 |
| 71752058 | -91.4458 |
| 3071746 | -91.3995 |
| 3612573 | -91.39 |
| 14687497 | -91.3619 |
| 7053486 | -91.3241 |
| 579398 | -91.302 |
| 74145 | -91.294 |
| 413671 | -91.2934 |
| 129008848 | -91.2802 |
| 38350655 | -91.2531 |
| 94940195 | -91.2425 |
| 82200638 | -91.2396 |
| 125416090 | -91.186 |
| 155888428 | -91.1275 |
| 12642748 | -91.1148 |
| 92214520 | -91.0706 |
| 10168 | -91.0682 |
| 71362251 | -91.0141 |
| 23696794 | -91.0044 |
| 87435342 | -90.9628 |
| 23661981 | -90.9618 |
| 23715428 | -90.9601 |
| 87318837 | -90.9573 |
| 24196322 | -90.9533 |
| 23666331 | -90.9508 |
| 3870576 | -90.9303 |
| 2847153 | -90.8574 |
| 2892875 | -90.8535 |
| 71325691 | -90.8448 |
| 86256957 | -90.8397 |
| 89677 | -90.7873 |
| 84372992 | -90.7737 |
| 57508711 | -90.7502 |
| 71425579 | -90.701 |
| 84373107 | -90.6834 |
| 3965218 | -90.6624 |
| 550738 | -90.6532 |
| 5228720 | -90.6457 |
| 5300192 | -90.6437 |
| 7460684 | -90.6404 |
| 71325063 | -90.6011 |
| 3840585 | -90.5836 |
| 3959118 | -90.5779 |
| 16568631 | -90.563 |
| 51537214 | -90.4882 |
| 14332561 | -90.4876 |
| 7343693 | -90.4743 |
| 7694413 | -90.3682 |
| 8551 | -90.2964 |
| 17297792 | -90.2503 |
| 71356913 | -90.2392 |
| 16205526 | -90.2349 |
| 56697362 | -90.2349 |
| 9797839 | -90.2074 |
| 7188003 | -90.1643 |
| 5300193 | -90.1556 |
| 412184 | -90.1064 |
| 92214519 | -90.0828 |
| 31477769 | -90.0603 |
| 129389 | -90.0539 |
| 6453094 | -90.0523 |
| 19154575 | -90.0455 |
| 71399358 | -90.0429 |
| 2749875 | -90.0117 |
| 7343695 | -89.9243 |
| 38357215 | -89.9239 |
| 21497700 | -89.9112 |
| 9019956 | -89.8328 |
| 4220639 | -89.8125 |
| 2917068 | -89.8067 |
| 135516791 | -89.7961 |
| 11507907 | -89.7959 |
| 46500497 | -89.7941 |
| 4243350 | -89.7858 |
| 3265888 | -89.7694 |
| 16416493 | -89.7686 |
| 4345463 | -89.765 |
| 617001 | -89.7516 |
| 7062419 | -89.7292 |
| 38357224 | -89.7142 |
| 626253 | -89.7126 |
| 193838 | -89.6985 |
| 5161690 | -89.6932 |
| 54332588 | -89.6837 |
| 13410197 | -89.6799 |
| 14139514 | -89.6745 |
| 19089713 | -89.6682 |
| 4401694 | -89.6637 |
| 13008123 | -89.6589 |
| 149693 | -89.6417 |
| 18925786 | -89.6259 |
| 4626007 | -89.6189 |
| 77725 | -89.6027 |
| 3352263 | -89.5932 |
| 3805465 | -89.552 |
| 84379165 | -89.5517 |
| 85967943 | -89.5508 |
| 412187 | -89.5474 |
| 84553260 | -89.5309 |
| 226106 | -89.5172 |
| 71394497 | -89.5121 |
| 412186 | -89.5091 |
| 331141 | -89.4717 |
| 6017159 | -89.4699 |
| 4614514 | -89.4585 |
| 40477923 | -89.4534 |
| 87728 | -89.4519 |
| 13510691 | -89.3437 |
| 5343613 | -89.3093 |
| 69607416 | -89.2944 |
| 18877552 | -89.2635 |
| 195788 | -89.26 |
| 2837135 | -89.2319 |
| 10685268 | -89.1441 |
| 14320105 | -89.1395 |
| 7154594 | -89.1348 |
| 42586887 | -89.099 |
| 53627553 | -89.0821 |
| 41149487 | -89.0187 |
| 3837515 | -89.0127 |
| 3331063 | -88.9834 |
| 12378369 | -88.9359 |
| 106049 | -88.8836 |
| 17587837 | -88.8727 |
| 112870 | -88.872 |
| 2837137 | -88.8706 |
| 20549 | -88.8603 |
| 2764162 | -88.8544 |
| 2847378 | -88.8542 |
| 7377778 | -88.8326 |
| 236918 | -88.8083 |
| 17354278 | -88.8063 |
| 3788004 | -88.7837 |
| 4381145 | -88.769 |
| 16205300 | -88.7648 |
| 14320106 | -88.7551 |
| 56481731 | -88.7349 |
| 21520014 | -88.7269 |
| 3538950 | -88.6811 |
| 3089796 | -88.6724 |
| 4206189 | -88.6705 |
| 110500478 | -88.6486 |
| 84379880 | -88.6423 |
| 5003831 | -88.6257 |
| 12966413 | -88.6076 |
| 71347905 | -88.588 |
| 138988056 | -88.5765 |
| 5875226 | -88.5193 |
| 14440510 | -88.4893 |
| 44288136 | -88.4761 |
| 30515146 | -88.4165 |
| 344313 | -88.3843 |
| 4338519 | -88.353 |
| 3830620 | -88.3485 |
| 3859780 | -88.3347 |
| 10802699 | -88.2907 |
| 4159121 | -88.2875 |
| 71445317 | -88.2827 |
| 2839153 | -88.2544 |
| 74146 | -88.2012 |
| 420666 | -88.1905 |
| 26435743 | -88.1755 |
| 7969622 | -88.175 |
| 133611900 | -88.1689 |
| 2847914 | -88.1565 |
| 84318781 | -88.1257 |
| 315790 | -88.1206 |
| 3090515 | -88.1179 |
| 57582998 | -88.0885 |
| 2933473 | -88.0445 |
| 21453711 | -88.0352 |
| 7344921 | -88.0165 |
| 2839444 | -88.0024 |
| 363065 | -88.0017 |
| 45359290 | -87.9998 |
| 222170 | -87.9831 |
| 15152091 | -87.9782 |
| 378693 | -87.9765 |
| 45034687 | -87.9703 |
| 94940061 | -87.9697 |
| 5387808 | -87.967 |
| 155011 | -87.9509 |
| 135719337 | -87.9436 |
| 361510 | -87.9337 |
| 7177252 | -87.9314 |
| 12318652 | -87.9136 |
| 5048656 | -87.8855 |
| 84378601 | -87.8457 |
| 631362 | -87.8419 |
| 2749864 | -87.8179 |
| 625242 | -87.8064 |
| 71399365 | -87.805 |
| 188678 | -87.8027 |
| 197351 | -87.7804 |
| 613895 | -87.7763 |
| 7863830 | -87.7758 |
| 101286218 | -87.767 |
| 44726113 | -87.7605 |
| 80834 | -87.7563 |
| 12336459 | -87.7339 |
| 16415109 | -87.7306 |
| 412183 | -87.7177 |
| 379539 | -87.7166 |
| 19154585 | -87.6952 |
| 5105399 | -87.6801 |
| 2748949 | -87.6736 |
| 6271568 | -87.6651 |
| 4415237 | -87.6548 |
| 71328559 | -87.6504 |
| 6104236 | -87.6499 |
| 23267431 | -87.6376 |
| 80851 | -87.6365 |
| 24721014 | -87.6288 |
| 71329023 | -87.6216 |
| 95528707 | -87.5999 |
| 86020143 | -87.5908 |
| 4406802 | -87.5886 |
| 4179245 | -87.574 |
| 6260036 | -87.5707 |
| 294388 | -87.5605 |
| 72747 | -87.5584 |
| 5326347 | -87.5548 |
| 7718568 | -87.5489 |
| 19792943 | -87.548 |
| 3126077 | -87.5094 |
| 5058246 | -87.5052 |
| 13906822 | -87.4914 |
| 13736891 | -87.447 |
| 147292 | -87.4392 |
| 71357489 | -87.4287 |
| 10615154 | -87.4251 |
| 15999227 | -87.419 |
| 71442175 | -87.4058 |
| 71414980 | -87.4038 |
| 3849711 | -87.3925 |
| 51537215 | -87.3841 |
| 5152048 | -87.3742 |
| 5490206 | -87.3738 |
| 147672 | -87.3733 |
| 3070902 | -87.3341 |
| 255082 | -87.3199 |
| 83523 | -87.3189 |
| 2886353 | -87.3017 |
| 335015 | -87.3002 |
| 11880227 | -87.2896 |
| 3154557 | -87.2473 |
| 19154558 | -87.2312 |
| 10947912 | -87.2212 |
| 2937895 | -87.2056 |
| 4035104 | -87.2051 |
| 197301 | -87.1927 |
| 3525837 | -87.1784 |
| 147671 | -87.1733 |
| 3090912 | -87.1605 |
| 134117733 | -87.1559 |
| 7079668 | -87.1542 |
| 6680 | -87.1533 |
| 11592174 | -87.1362 |
| 125416091 | -87.13 |
| 52100662 | -87.11 |
| 71413309 | -87.1037 |
| 3280108 | -87.0992 |
| 20194310 | -87.0867 |
| 80745 | -87.0817 |
| 2892372 | -87.0582 |
| 331628 | -87.0578 |
| 511525 | -87.0547 |
| 3090514 | -87.0373 |
| 11110093 | -87.0257 |
| 5054260 | -87.0239 |
| 58321336 | -87.0213 |
| 2933120 | -87.0148 |
| 84553428 | -87.0142 |
| 31477775 | -87.0105 |
| 7060274 | -86.9959 |
| 46500766 | -86.9771 |
| 20194335 | -86.9706 |
| 71343342 | -86.9644 |
| 41673495 | -86.9634 |
| 44784325 | -86.9549 |
| 25905 | -86.9547 |
| 137163265 | -86.8848 |
| 151995 | -86.8813 |
| 13162745 | -86.8674 |
| 13605997 | -86.8573 |
| 31413 | -86.8482 |
| 304458 | -86.841 |
| 49696985 | -86.8356 |
| 622339 | -86.8109 |
| 11779662 | -86.7946 |
| 71325694 | -86.7772 |
| 82200640 | -86.7749 |
| 3887945 | -86.7738 |
| 18877549 | -86.7737 |
| 84373096 | -86.7725 |
| 2847623 | -86.771 |
| 4129741 | -86.7508 |
| 71353498 | -86.75 |
| 2748415 | -86.7476 |
| 7095967 | -86.7456 |
| 2848383 | -86.7216 |
| 71319036 | -86.7137 |
| 102955 | -86.7001 |
| 71365423 | -86.698 |
| 84373070 | -86.6939 |
| 12324276 | -86.6813 |
| 90269327 | -86.6467 |
| 426725 | -86.6417 |
| 88253 | -86.6364 |
| 3126044 | -86.6306 |
| 3009664 | -86.6257 |
| 12182525 | -86.6219 |
| 439020 | -86.62 |
| 108796815 | -86.6145 |
| 71399364 | -86.6139 |
| 7072973 | -86.5941 |
| 12313085 | -86.5934 |
| 3754993 | -86.5834 |
| 28531 | -86.5529 |
| 10358030 | -86.5482 |
| 12324271 | -86.535 |
| 262099 | -86.5287 |
| 4613044 | -86.5284 |
| 7153805 | -86.4922 |
| 12490500 | -86.4893 |
| 2898767 | -86.4892 |
| 46498897 | -86.4888 |
| 92651860 | -86.4613 |
| 57958 | -86.4527 |
| 147669 | -86.4432 |
| 625632 | -86.4396 |
| 14379528 | -86.434 |
| 335705 | -86.4157 |
| 2829333 | -86.4133 |
| 85528929 | -86.4104 |
| 20432761 | -86.3968 |
| 3320955 | -86.388 |
| 66479 | -86.3571 |
| 17263904 | -86.3465 |
| 122707271 | -86.3454 |
| 2933860 | -86.3304 |
| 5139379 | -86.3115 |
| 20414990 | -86.3069 |
| 374507 | -86.2759 |
| 9581276 | -86.2721 |
| 3108086 | -86.2671 |
| 23276522 | -86.2621 |
| 7103642 | -86.2412 |
| 4598057 | -86.239 |
| 69057 | -86.2367 |
| 5061436 | -86.2267 |
| 122902 | -86.1936 |
| 5205317 | -86.1909 |
| 4037606 | -86.1745 |
| 147665 | -86.1432 |
| 67011 | -86.143 |
| 2856876 | -86.1099 |
| 5320212 | -86.1042 |
| 90659 | -86.1035 |
| 2930712 | -86.0993 |
| 283899 | -86.0806 |
| 10757461 | -86.0778 |
| 46500767 | -86.0728 |
| 7114955 | -86.0713 |
| 2838304 | -86.0683 |
| 331627 | -86.0661 |
| 71399359 | -86.0634 |
| 235843 | -86.0504 |
| 2835180 | -86.0495 |
| 5048591 | -86.0439 |
| 11035197 | -86.0211 |
| 337182 | -85.9998 |
| 331638 | -85.9996 |
| 2892351 | -85.9761 |
| 147675 | -85.9757 |
| 2847852 | -85.9685 |
| 16659185 | -85.9576 |
| 28476370 | -85.9529 |
| 84320699 | -85.9468 |
| 5171297 | -85.9375 |
| 77118 | -85.9364 |
| 506610 | -85.9328 |
| 67195 | -85.9292 |
| 13116586 | -85.9147 |
| 3754690 | -85.901 |
| 3561972 | -85.8931 |
| 354963 | -85.8901 |
| 67008 | -85.8871 |
| 423406 | -85.8726 |
| 95987981 | -85.8586 |
| 7074403 | -85.8565 |
| 25082968 | -85.8532 |
| 84373000 | -85.8495 |
| 87406 | -85.84 |
| 570342 | -85.837 |
| 619349 | -85.8323 |
| 23662526 | -85.8271 |
| 12780398 | -85.8219 |
| 14595174 | -85.7903 |
| 359018 | -85.7888 |
| 71341369 | -85.7768 |
| 2892156 | -85.7751 |
| 14816593 | -85.7745 |
| 7969695 | -85.7709 |
| 7064882 | -85.7395 |
| 3743766 | -85.7373 |
| 2749314 | -85.7119 |
| 160476 | -85.6874 |
| 139035518 | -85.67 |
| 3147193 | -85.6656 |
| 4443133 | -85.665 |
| 13380585 | -85.6617 |
| 6111772 | -85.6604 |
| 72739 | -85.6349 |
| 84373090 | -85.63 |
| 132248 | -85.6246 |
| 314075 | -85.6239 |
| 20485972 | -85.6189 |
| 627321 | -85.6179 |
| 71426393 | -85.5935 |
| 13970499 | -85.5817 |
| 2838584 | -85.575 |
| 131350 | -85.5686 |
| 16952536 | -85.5681 |
| 38357209 | -85.5623 |
| 54183254 | -85.562 |
| 11142042 | -85.5547 |
| 7053214 | -85.5465 |
| 8235936 | -85.5449 |
| 12256454 | -85.5406 |
| 9635553 | -85.5157 |
| 331626 | -85.5079 |
| 3778978 | -85.5058 |
| 71415670 | -85.4804 |
| 13328362 | -85.4775 |
| 24319526 | -85.4668 |
| 3135548 | -85.4612 |
| 3325451 | -85.4599 |
| 90471648 | -85.4579 |
| 3757176 | -85.4374 |
| 24319544 | -85.4353 |
| 2893068 | -85.4341 |
| 20366329 | -85.428 |
| 98128054 | -85.4082 |
| 22696541 | -85.3956 |
| 323139 | -85.3953 |
| 613892 | -85.3889 |
| 626899 | -85.3886 |
| 3113290 | -85.3712 |
| 624966 | -85.3509 |
| 156677 | -85.3487 |
| 630572 | -85.3414 |
| 630215 | -85.3392 |
| 33784 | -85.3382 |
| 9994012 | -85.3334 |
| 44287920 | -85.3193 |
| 91664854 | -85.3182 |
| 17283767 | -85.3065 |
| 3100810 | -85.3048 |
| 15620928 | -85.3042 |
| 71329481 | -85.2919 |
| 8811074 | -85.2883 |
| 71399362 | -85.2804 |
| 135462137 | -85.2686 |
| 12318648 | -85.2652 |
| 135404103 | -85.263 |
| 82200639 | -85.257 |
| 11652569 | -85.2552 |
| 5174766 | -85.239 |
| 71412296 | -85.2372 |
| 14534995 | -85.2322 |
| 13256304 | -85.225 |
| 138112320 | -85.2112 |
| 13214483 | -85.2101 |
| 303624 | -85.2095 |
| 3564104 | -85.2005 |
| 94940194 | -85.1906 |
| 5920527 | -85.1799 |
| 8867476 | -85.1766 |
| 149895 | -85.1716 |
| 84376379 | -85.1672 |
| 131861851 | -85.145 |
| 91665607 | -85.1433 |
| 17144610 | -85.1361 |
| 46949535 | -85.1361 |
| 3126053 | -85.1317 |
| 16068 | -85.1302 |
| 10870976 | -85.1085 |
| 16800486 | -85.1068 |
| 3126048 | -85.1013 |
| 92237096 | -85.0581 |
| 13423601 | -85.0544 |
| 222227 | -85.0535 |
| 22062352 | -85.0518 |
| 4733 | -85.046 |
| 458834 | -85.0426 |
| 424403 | -85.0289 |
| 71347012 | -85.015 |
| 259861 | -85.0017 |
| 7969636 | -84.9999 |
| 4663427 | -84.9766 |
| 443807 | -84.9644 |
| 3861237 | -84.9564 |
| 60020227 | -84.9481 |
| 16205613 | -84.9417 |
| 4967416 | -84.9394 |
| 3764511 | -84.9376 |
| 13134882 | -84.9298 |
| 11880225 | -84.9205 |
| 15483121 | -84.9142 |
| 4964468 | -84.9129 |
| 317256 | -84.8984 |
| 71349161 | -84.897 |
| 3155948 | -84.8717 |
| 16762955 | -84.8688 |
| 2749495 | -84.8548 |
| 6062206 | -84.8334 |
| 12324272 | -84.833 |
| 4462908 | -84.8295 |
| 2832164 | -84.8267 |
| 71319042 | -84.8258 |
| 91894 | -84.8094 |
| 12444971 | -84.7976 |
| 8219072 | -84.7911 |
| 149791 | -84.7866 |
| 23667353 | -84.7672 |
| 87523472 | -84.761 |
| 71319064 | -84.7609 |
| 24181116 | -84.7596 |
| 50434 | -84.7497 |
| 42559548 | -84.7367 |
| 630908 | -84.7225 |
| 12230782 | -84.7173 |
| 7083422 | -84.7155 |
| 280737 | -84.7055 |
| 7331339 | -84.7027 |
| 44251286 | -84.6959 |
| 71371853 | -84.6957 |
| 11782400 | -84.6892 |
| 7122777 | -84.6711 |
| 4497953 | -84.6649 |
| 11834180 | -84.6619 |
| 2752531 | -84.657 |
| 3668957 | -84.6545 |
| 20317508 | -84.6503 |
| 71329750 | -84.6496 |
| 13955348 | -84.6311 |
| 318176 | -84.6297 |
| 21062 | -84.6283 |
| 7064918 | -84.6147 |
| 7230010 | -84.6116 |
| 4553296 | -84.6057 |
| 7083423 | -84.6044 |
| 18877555 | -84.6013 |
| 71319070 | -84.6009 |
| 291740 | -84.5999 |
| 631009 | -84.5984 |
| 396203 | -84.5751 |
| 2828783 | -84.5594 |
| 3103608 | -84.5589 |
| 91664802 | -84.5578 |
| 46782753 | -84.5416 |
| 622513 | -84.5364 |
| 11185595 | -84.5111 |
| 20414995 | -84.51 |
| 10403632 | -84.499 |
| 182190 | -84.4825 |
| 3090508 | -84.48 |
| 12360344 | -84.4771 |
| 10639 | -84.4738 |
| 30515338 | -84.4676 |
| 7079669 | -84.4531 |
| 91664798 | -84.4497 |
| 254705 | -84.449 |
| 14250673 | -84.4316 |
| 90673963 | -84.4158 |
| 7730276 | -84.408 |
| 3550312 | -84.3913 |
| 89063 | -84.3871 |
| 92651859 | -84.3849 |
| 113243 | -84.3677 |
| 619461 | -84.3661 |
| 443808 | -84.3475 |
| 50918921 | -84.3456 |
| 5235039 | -84.3417 |
| 56632138 | -84.3283 |
| 3100911 | -84.3276 |
| 84375473 | -84.3269 |
| 15592816 | -84.3265 |
| 225598 | -84.3078 |
| 372654 | -84.3073 |
| 28476461 | -84.2944 |
| 423402 | -84.2857 |
| 127257877 | -84.2781 |
| 15778586 | -84.271 |
| 14447720 | -84.2524 |
| 3929987 | -84.2452 |
| 626912 | -84.237 |
| 3025849 | -84.2294 |
| 71325062 | -84.2133 |
| 2749867 | -84.2128 |
| 7421451 | -84.2114 |
| 84554417 | -84.2062 |
| 270486 | -84.2027 |
| 90473387 | -84.1952 |
| 56839171 | -84.1949 |
| 374058 | -84.1946 |
| 3678450 | -84.1923 |
| 3721458 | -84.192 |
| 12069476 | -84.1871 |
| 628974 | -84.1823 |
| 10958178 | -84.1808 |
| 423403 | -84.1561 |
| 73995918 | -84.1545 |
| 17586893 | -84.1502 |
| 16880175 | -84.147 |
| 12318647 | -84.1433 |
| 85540 | -84.1283 |
| 71597362 | -84.1273 |
| 76264 | -84.1167 |
| 13675111 | -84.1131 |
| 71353499 | -84.1125 |
| 3418526 | -84.1111 |
| 4427889 | -84.1095 |
| 3083633 | -84.102 |
| 21453696 | -84.0709 |
| 44150898 | -84.0703 |
| 629326 | -84.0682 |
| 71405345 | -84.0669 |
| 12357435 | -84.0653 |
| 3126058 | -84.0636 |
| 8901891 | -84.0582 |
| 459276 | -84.0563 |
| 559503 | -84.039 |
| 11098600 | -84.0253 |
| 98128057 | -84.0096 |
| 8142815 | -84.0037 |
| 12342683 | -84.0031 |
| 10086148 | -83.9886 |
| 261069 | -83.977 |
| 23663644 | -83.9745 |
| 7694275 | -83.9702 |
| 337183 | -83.9676 |
| 50884020 | -83.9663 |
| 443801 | -83.9656 |
| 3625488 | -83.963 |
| 624085 | -83.939 |
| 16954045 | -83.9341 |
| 122751 | -83.9273 |
| 629785 | -83.9253 |
| 3124730 | -83.9199 |
| 3125184 | -83.9068 |
| 432252 | -83.9014 |
| 13599687 | -83.901 |
| 5319342 | -83.8964 |
| 7226818 | -83.8827 |
| 11098986 | -83.8817 |
| 44287910 | -83.8768 |
| 3469323 | -83.8673 |
| 126037 | -83.8558 |
| 98115208 | -83.8536 |
| 5704845 | -83.8513 |
| 13788856 | -83.851 |
| 3905783 | -83.8509 |
| 160712 | -83.8484 |
| 5048592 | -83.8437 |
| 72752 | -83.8403 |
| 12416404 | -83.8253 |
| 124926260 | -83.8243 |
| 18877560 | -83.8178 |
| 71410411 | -83.8094 |
| 71374597 | -83.8046 |
| 20414989 | -83.7968 |
| 76647 | -83.7845 |
| 10903651 | -83.7826 |
| 195786 | -83.7806 |
| 3678893 | -83.7708 |
| 47752 | -83.766 |
| 71381687 | -83.7652 |
| 331633 | -83.7618 |
| 344310 | -83.7597 |
| 80833 | -83.7513 |
| 56445 | -83.7272 |
| 11066989 | -83.7264 |
| 7980504 | -83.71 |
| 4395272 | -83.7076 |
| 10040432 | -83.7 |
| 11077433 | -83.6907 |
| 84320710 | -83.6868 |
| 23051730 | -83.6867 |
| 66591565 | -83.6848 |
| 12865600 | -83.6832 |
| 159926 | -83.68 |
| 236917 | -83.6792 |
| 3089800 | -83.6765 |
| 620551 | -83.664 |
| 9581278 | -83.6568 |
| 3126029 | -83.6465 |
| 12756410 | -83.6347 |
| 355595 | -83.6338 |
| 84379319 | -83.6231 |
| 44575883 | -83.5912 |
| 5704843 | -83.5751 |
| 84376221 | -83.575 |
| 40468805 | -83.5576 |
| 16206120 | -83.5567 |
| 4165774 | -83.5432 |
| 133505 | -83.5411 |
| 12509074 | -83.5169 |
| 156465 | -83.5016 |
| 124430675 | -83.4953 |
| 53440214 | -83.4777 |
| 374505 | -83.476 |
| 13482589 | -83.4677 |
| 51056136 | -83.461 |
| 11573214 | -83.4596 |
| 278670 | -83.4581 |
| 42647718 | -83.4557 |
| 42559549 | -83.4494 |
| 4533665 | -83.443 |
| 82199246 | -83.4424 |
| 3149913 | -83.4397 |
| 13005613 | -83.438 |
| 12808260 | -83.4291 |
| 2749868 | -83.4223 |
| 5326346 | -83.4211 |
| 90569 | -83.4177 |
| 15965431 | -83.4121 |
| 4676200 | -83.4071 |
| 2930695 | -83.3979 |
| 56697077 | -83.388 |
| 2749877 | -83.3769 |
| 11727234 | -83.3672 |
| 12325134 | -83.3517 |
| 131845149 | -83.3508 |
| 54224488 | -83.3253 |
| 144045074 | -83.3084 |
| 7969789 | -83.2882 |
| 71334793 | -83.2836 |
| 3102924 | -83.2796 |
| 71437089 | -83.2791 |
| 14563137 | -83.2777 |
| 4988403 | -83.2703 |
| 361513 | -83.2602 |
| 76267 | -83.252 |
| 2892157 | -83.248 |
| 71407403 | -83.2297 |
| 84378381 | -83.225 |
| 10870113 | -83.2191 |
| 126726 | -83.2148 |
| 84379968 | -83.2036 |
| 18877556 | -83.2001 |
| 26343 | -83.1711 |
| 16205562 | -83.168 |
| 16205903 | -83.1542 |
| 93949197 | -83.1454 |
| 13710020 | -83.1441 |
| 71329548 | -83.1344 |
| 3124731 | -83.1284 |
| 7104294 | -83.1201 |
| 10085264 | -83.1159 |
| 16421299 | -83.1042 |
| 16880197 | -83.0997 |
| 8960 | -83.0936 |
| 92221389 | -83.0901 |
| 44287706 | -83.0898 |
| 3271702 | -83.0808 |
| 44251291 | -83.0687 |
| 8235946 | -83.0686 |
| 3106278 | -83.0654 |
| 20456767 | -83.0507 |
| 3831797 | -83.0462 |
| 3143428 | -83.0107 |
| 78175850 | -83.0101 |
| 124430676 | -83.0086 |
| 3125602 | -83.0084 |
| 363264 | -82.9908 |
| 159258 | -82.9861 |
| 127173 | -82.9817 |
| 82030677 | -82.969 |
| 53297773 | -82.9682 |
| 71345801 | -82.9662 |
| 44721070 | -82.9504 |
| 5125574 | -82.9469 |
| 624945 | -82.9466 |
| 3125609 | -82.9465 |
| 53440301 | -82.9461 |
| 71446055 | -82.9422 |
| 2856897 | -82.9214 |
| 53440271 | -82.9099 |
| 42647717 | -82.9064 |
| 11833329 | -82.9003 |
| 354953 | -82.8921 |
| 344340 | -82.8761 |
| 7969792 | -82.8719 |
| 173905 | -82.8614 |
| 384115 | -82.8422 |
| 92252410 | -82.8383 |
| 9817337 | -82.8354 |
| 28476886 | -82.813 |
| 84378379 | -82.8058 |
| 13181158 | -82.7986 |
| 10403757 | -82.7955 |
| 45043556 | -82.7945 |
| 49695419 | -82.7785 |
| 17144608 | -82.7767 |
| 56439 | -82.7668 |
| 5972358 | -82.7634 |
| 4054635 | -82.7568 |
| 8030558 | -82.7553 |
| 2847035 | -82.7538 |
| 10778954 | -82.7456 |
| 4267041 | -82.7443 |
| 624934 | -82.7413 |
| 131859082 | -82.7407 |
| 3967018 | -82.7347 |
| 14688012 | -82.7304 |
| 12220565 | -82.7283 |
| 354143 | -82.7265 |
| 625092 | -82.7197 |
| 139270371 | -82.7185 |
| 49694130 | -82.7027 |
| 12003337 | -82.6982 |
| 3662101 | -82.6965 |
| 7064883 | -82.6611 |
| 371563 | -82.6475 |
| 15559229 | -82.647 |
| 5128995 | -82.6434 |
| 14039099 | -82.6424 |
| 88001 | -82.6406 |
| 3089795 | -82.6367 |
| 8901890 | -82.6244 |
| 21637461 | -82.6208 |
| 354136 | -82.6183 |
| 10066319 | -82.6068 |
| 44287645 | -82.5946 |
| 110499464 | -82.5896 |
| 2833771 | -82.576 |
| 71440549 | -82.5677 |
| 91362 | -82.5641 |
| 261068 | -82.5593 |
| 7390523 | -82.5334 |
| 4158921 | -82.5285 |
| 620365 | -82.5155 |
| 350511 | -82.4888 |
| 431929 | -82.4888 |
| 367128 | -82.4853 |
| 52493 | -82.4836 |
| 88430 | -82.4799 |
| 7053489 | -82.4796 |
| 4345775 | -82.4745 |
| 3929988 | -82.4611 |
| 21572844 | -82.4584 |
| 13539447 | -82.4565 |
| 13439293 | -82.4533 |
| 13380575 | -82.4482 |
| 2754660 | -82.4332 |
| 345814 | -82.4318 |
| 54412600 | -82.4239 |
| 616266 | -82.4217 |
| 56691480 | -82.408 |
| 5319801 | -82.401 |
| 613891 | -82.3995 |
| 32209 | -82.3892 |
| 10517868 | -82.382 |
| 133641338 | -82.3729 |
| 16764465 | -82.3623 |
| 23260727 | -82.362 |
| 71404343 | -82.3586 |
| 10017076 | -82.3539 |
| 2749602 | -82.3533 |
| 96002062 | -82.3519 |
| 5021913 | -82.3486 |
| 133506 | -82.3368 |
| 71399363 | -82.3336 |
| 11529868 | -82.3209 |
| 71399354 | -82.3152 |
| 95987982 | -82.3147 |
| 92284877 | -82.3083 |
| 70518802 | -82.2918 |
| 73490 | -82.2906 |
| 271591 | -82.2884 |
| 7721316 | -82.2877 |
| 53440421 | -82.276 |
| 11482604 | -82.2754 |
| 9883304 | -82.2712 |
| 236912 | -82.2512 |
| 4114834 | -82.2487 |
| 7062420 | -82.2461 |
| 7392102 | -82.2454 |
| 5928598 | -82.2323 |
| 2917207 | -82.2313 |
| 482557 | -82.2183 |
| 12865589 | -82.2142 |
| 167525 | -82.2095 |
| 381097 | -82.1982 |
| 3463237 | -82.1943 |
| 71410410 | -82.1919 |
| 3428212 | -82.1849 |
| 71319033 | -82.1844 |
| 146101 | -82.1803 |
| 12334640 | -82.1796 |
| 13412786 | -82.1745 |
| 31534 | -82.1717 |
| 101317808 | -82.1575 |
| 620322 | -82.1505 |
| 344129 | -82.1493 |
| 3126050 | -82.1403 |
| 94938900 | -82.1249 |
| 71319041 | -82.12 |
| 16954063 | -82.1146 |
| 71327785 | -82.1129 |
| 3100914 | -82.1066 |
| 337849 | -82.1058 |
| 71402041 | -82.1041 |
| 12812102 | -82.1017 |
| 4589998 | -82.0884 |
| 11255215 | -82.0876 |
| 17191 | -82.0839 |
| 86109348 | -82.0831 |
| 638569 | -82.0769 |
| 18526640 | -82.0721 |
| 418574 | -82.0715 |
| 6455094 | -82.0709 |
| 88436 | -82.0608 |
| 3146261 | -82.0557 |
| 13455966 | -82.0552 |
| 350271 | -82.0538 |
| 70669 | -82.0449 |
| 158681 | -82.041 |
| 2833773 | -82.036 |
| 270752 | -82.0224 |
| 97691 | -82.0173 |
| 44917092 | -82.0136 |
| 3094361 | -82.0105 |
| 3715877 | -82.0094 |
| 4429207 | -82.0034 |
| 14274943 | -81.9997 |
| 12548 | -81.9975 |
| 3600317 | -81.9916 |
| 442765 | -81.9661 |
| 76831 | -81.9656 |
| 57357605 | -81.9617 |
| 13317888 | -81.961 |
| 5333690 | -81.9521 |
| 20414994 | -81.9478 |
| 3410999 | -81.9448 |
| 12180561 | -81.9426 |
| 3091454 | -81.9323 |
| 20414997 | -81.9247 |
| 26435742 | -81.9219 |
| 3020434 | -81.9195 |
| 2892180 | -81.9089 |
| 110500517 | -81.8958 |
| 92284791 | -81.8924 |
| 40503 | -81.8892 |
| 77114 | -81.8883 |
| 135416853 | -81.8848 |
| 5339597 | -81.8778 |
| 112757967 | -81.8752 |
| 16205362 | -81.872 |
| 3090915 | -81.8536 |
| 15937278 | -81.8488 |
| 98121235 | -81.8433 |
| 6108061 | -81.8431 |
| 3746625 | -81.8411 |
| 3090913 | -81.833 |
| 67203912 | -81.8288 |
| 71446056 | -81.8139 |
| 3453577 | -81.8054 |
| 5013064 | -81.7848 |
| 95725 | -81.7814 |
| 13162744 | -81.7778 |
| 23349628 | -81.7769 |
| 147673 | -81.7696 |
| 363869 | -81.7634 |
| 5150597 | -81.7608 |
| 135844 | -81.7348 |
| 7062656 | -81.7342 |
| 3123178 | -81.7312 |
| 86142772 | -81.7248 |
| 15665217 | -81.7244 |
| 344317 | -81.7145 |
| 109836 | -81.7075 |
| 132243 | -81.7016 |
| 2752532 | -81.6978 |
| 619609 | -81.6908 |
| 18870669 | -81.6904 |
| 16952511 | -81.6897 |
| 3418613 | -81.6828 |
| 15743473 | -81.6797 |
| 71410869 | -81.6708 |
| 73996030 | -81.6671 |
| 4658671 | -81.6647 |
| 84555133 | -81.6645 |
| 41149577 | -81.6477 |
| 24241 | -81.6403 |
| 60208839 | -81.6403 |
| 10427286 | -81.6294 |
| 10125866 | -81.6235 |
| 40504 | -81.622 |
| 3142375 | -81.6209 |
| 22643726 | -81.5997 |
| 22956534 | -81.5885 |
| 5704919 | -81.5868 |
| 23051731 | -81.5731 |
| 2871272 | -81.5628 |
| 22643714 | -81.5596 |
| 318390 | -81.5393 |
| 442750 | -81.5331 |
| 20494165 | -81.5323 |
| 10518158 | -81.5293 |
| 155380 | -81.5291 |
| 28846569 | -81.5192 |
| 22829993 | -81.4822 |
| 57352734 | -81.4807 |
| 22959934 | -81.479 |
| 57352733 | -81.474 |
| 2847065 | -81.4618 |
| 625088 | -81.4534 |
| 92210352 | -81.4469 |
| 4220118 | -81.443 |
| 423405 | -81.4387 |
| 344321 | -81.4332 |
| 14320098 | -81.4323 |
| 5124769 | -81.4211 |
| 281889 | -81.4166 |
| 71361559 | -81.4152 |
| 7097710 | -81.4111 |
| 96469022 | -81.3865 |
| 2890433 | -81.386 |
| 442729 | -81.3794 |
| 71442525 | -81.3763 |
| 5326348 | -81.3649 |
| 131667939 | -81.3623 |
| 182301 | -81.3621 |
| 42586886 | -81.3575 |
| 3321713 | -81.3557 |
| 11231645 | -81.3459 |
| 6454002 | -81.3385 |
| 5022029 | -81.326 |
| 7164286 | -81.3231 |
| 7115189 | -81.3078 |
| 7115187 | -81.3039 |
| 291367 | -81.2921 |
| 86142836 | -81.2911 |
| 19705590 | -81.2898 |
| 149782 | -81.2882 |
| 337848 | -81.2871 |
| 28477031 | -81.2782 |
| 71329157 | -81.2728 |
| 442766 | -81.2653 |
| 146022354 | -81.2558 |
| 4153655 | -81.2481 |
| 67340699 | -81.2358 |
| 443789 | -81.2342 |
| 99188 | -81.2308 |
| 335014 | -81.2208 |
| 16528 | -81.2162 |
| 53437513 | -81.2155 |
| 222169 | -81.2132 |
| 18877557 | -81.2037 |
| 59279133 | -81.1979 |
| 7179225 | -81.1976 |
| 85781266 | -81.1947 |
| 51138600 | -81.1921 |
| 361512 | -81.1892 |
| 164046 | -81.1841 |
| 46500437 | -81.1783 |
| 16762870 | -81.175 |
| 331630 | -81.1743 |
| 13270885 | -81.161 |
| 87342 | -81.1583 |
| 2837521 | -81.1558 |
| 28846498 | -81.1558 |
| 72742 | -81.1507 |
| 20240722 | -81.1503 |
| 235734 | -81.1461 |
| 4175705 | -81.1451 |
| 331635 | -81.1208 |
| 92297216 | -81.1194 |
| 71401825 | -81.0989 |
| 82199245 | -81.0953 |
| 71371928 | -81.092 |
| 2892261 | -81.0903 |
| 12309204 | -81.0712 |
| 104186 | -81.0688 |
| 12383760 | -81.0662 |
| 363709 | -81.0629 |
| 313995 | -81.0594 |
| 125152 | -81.0576 |
| 342672 | -81.056 |
| 7529565 | -81.0555 |
| 40468806 | -81.0516 |
| 23623134 | -81.0474 |
| 3908369 | -81.0416 |
| 624824 | -81.033 |
| 109809 | -81.0307 |
| 100214 | -81.0248 |
| 13412787 | -81.0168 |
| 40553271 | -81.0125 |
| 40578925 | -81.0073 |
| 13007500 | -81.0054 |
| 12314923 | -80.989 |
| 3793750 | -80.985 |
| 630594 | -80.9776 |
| 7065339 | -80.9756 |
| 10379693 | -80.9609 |
| 13013280 | -80.9414 |
| 87014 | -80.9365 |
| 315276 | -80.9227 |
| 4626945 | -80.9052 |
| 86012754 | -80.8981 |
| 4357194 | -80.8916 |
| 54462239 | -80.8857 |
| 3108087 | -80.8836 |
| 14419207 | -80.8773 |
| 13766143 | -80.8758 |
| 4376010 | -80.8717 |
| 92255351 | -80.8686 |
| 16203753 | -80.8467 |
| 631405 | -80.8343 |
| 3013870 | -80.823 |
| 14239297 | -80.8173 |
| 7420410 | -80.8121 |
| 442756 | -80.8115 |
| 3483941 | -80.8106 |
| 373013 | -80.8066 |
| 277844 | -80.799 |
| 71335301 | -80.7966 |
| 21762900 | -80.7949 |
| 3040635 | -80.7944 |
| 56693988 | -80.7897 |
| 71361131 | -80.7881 |
| 12313148 | -80.7864 |
| 71361558 | -80.7864 |
| 7062657 | -80.7726 |
| 232873 | -80.7605 |
| 71319038 | -80.7596 |
| 337812 | -80.7565 |
| 10686237 | -80.7521 |
| 152706 | -80.742 |
| 396202 | -80.7349 |
| 350390 | -80.7307 |
| 2916691 | -80.7037 |
| 11674044 | -80.7015 |
| 7063302 | -80.6931 |
| 71365020 | -80.6906 |
| 160887 | -80.6826 |
| 357501 | -80.6795 |
| 86877736 | -80.6781 |
| 147621 | -80.6692 |
| 2911965 | -80.6689 |
| 10336479 | -80.6673 |
| 71319069 | -80.6642 |
| 390050 | -80.6632 |
| 7179561 | -80.6627 |
| 236913 | -80.6623 |
| 11808929 | -80.6572 |
| 2847134 | -80.6509 |
| 46494782 | -80.6455 |
| 15259336 | -80.641 |
| 12488554 | -80.6333 |
| 10163 | -80.6321 |
| 354134 | -80.6303 |
| 3702732 | -80.6303 |
| 13681829 | -80.6293 |
| 71399355 | -80.6242 |
| 3125601 | -80.6219 |
| 12631819 | -80.616 |
| 355494 | -80.614 |
| 3335146 | -80.6084 |
| 71410866 | -80.6008 |
| 71379712 | -80.6 |
| 3770675 | -80.5983 |
| 20419 | -80.5959 |
| 10539812 | -80.5918 |
| 93926 | -80.5796 |
| 4217712 | -80.5754 |
| 379538 | -80.5676 |
| 44149742 | -80.5566 |
| 625611 | -80.5514 |
| 91674734 | -80.5508 |
| 559564 | -80.5494 |
| 71324948 | -80.5467 |
| 92255376 | -80.5464 |
| 84375447 | -80.5456 |
| 625633 | -80.5436 |
| 5384873 | -80.5404 |
| 82200939 | -80.538 |
| 177458 | -80.5355 |
| 90953 | -80.5341 |
| 2835177 | -80.5329 |
| 124464350 | -80.5315 |
| 626653 | -80.5296 |
| 4544448 | -80.5256 |
| 110499500 | -80.5231 |
| 15743471 | -80.5207 |
| 4593548 | -80.5205 |
| 146022355 | -80.5016 |
| 21918414 | -80.4981 |
| 4027225 | -80.4976 |
| 617790 | -80.4936 |
| 11543612 | -80.4805 |
| 94940060 | -80.48 |
| 278318 | -80.4787 |
| 23260551 | -80.4783 |
| 71313110 | -80.4755 |
| 23260550 | -80.4743 |
| 15945181 | -80.4641 |
| 23617 | -80.4632 |
| 103229 | -80.4523 |
| 354135 | -80.4389 |
| 12005091 | -80.4338 |
| 3089797 | -80.4316 |
| 184395 | -80.4285 |
| 5011815 | -80.3932 |
| 28578 | -80.3767 |
| 44251212 | -80.3719 |
| 2846999 | -80.3687 |
| 71365497 | -80.3613 |
| 624935 | -80.3584 |
| 5156062 | -80.3519 |
| 9636987 | -80.3509 |
| 56438 | -80.3418 |
| 10207 | -80.3337 |
| 12443362 | -80.3247 |
| 4189238 | -80.3157 |
| 433411 | -80.3152 |
| 2932129 | -80.3124 |
| 341033 | -80.3101 |
| 95707 | -80.3081 |
| 4269884 | -80.3055 |
| 56689666 | -80.2986 |
| 10469084 | -80.2984 |
| 232527 | -80.2886 |
| 433917 | -80.273 |
| 92214784 | -80.2693 |
| 540862 | -80.2615 |
| 5319503 | -80.2408 |
| 21239136 | -80.2399 |
| 3100887 | -80.2209 |
| 7077528 | -80.2202 |
| 14094535 | -80.2195 |
| 5142634 | -80.2194 |
| 13331055 | -80.2175 |
| 636723 | -80.216 |
| 117101 | -80.2048 |
| 71402200 | -80.1958 |
| 44287918 | -80.1874 |
| 132244 | -80.1866 |
| 3126042 | -80.1852 |
| 11771669 | -80.1783 |
| 426250 | -80.1782 |
| 3036666 | -80.1782 |
| 67208 | -80.1753 |
| 282271 | -80.1708 |
| 4585422 | -80.1562 |
| 22673158 | -80.1463 |
| 374059 | -80.1393 |
| 628034 | -80.1241 |
| 71319063 | -80.1223 |
| 6282947 | -80.1196 |
| 2749601 | -80.1171 |
| 151858 | -80.1164 |
| 624579 | -80.1079 |
| 12386943 | -80.0986 |
| 71415473 | -80.0782 |
| 44145850 | -80.0764 |
| 624954 | -80.0744 |
| 7063303 | -80.066 |
| 630348 | -80.0605 |
| 92301622 | -80.0542 |
| 395766 | -80.0349 |
| 3808589 | -80.0307 |
| 42586889 | -80.0187 |
| 335706 | -80.0137 |
| 19685744 | -80.0088 |
| 11770969 | -80.0076 |
| 622752 | -80.0029 |
| 56687832 | -79.9969 |
| 361515 | -79.9895 |
| 5151543 | -79.9865 |
| 12424509 | -79.9813 |
| 85822847 | -79.9803 |
| 4191680 | -79.9692 |
| 57351861 | -79.969 |
| 14375937 | -79.9678 |
| 16762569 | -79.9606 |
| 4228367 | -79.9567 |
| 41929 | -79.9529 |
| 12388002 | -79.9419 |
| 6032119 | -79.9412 |
| 71329158 | -79.9377 |
| 7420408 | -79.9339 |
| 92258678 | -79.9316 |
| 13190720 | -79.9314 |
| 3664899 | -79.9257 |
| 71384163 | -79.9101 |
| 151669 | -79.91 |
| 5743850 | -79.9077 |
| 4917450 | -79.8965 |
| 622160 | -79.8931 |
| 4649699 | -79.8896 |
| 20414993 | -79.88 |
| 7330511 | -79.878 |
| 14767281 | -79.8779 |
| 10936407 | -79.877 |
| 3261304 | -79.872 |
| 41934 | -79.865 |
| 13708175 | -79.8582 |
| 66530 | -79.851 |
| 86021351 | -79.8419 |
| 5076150 | -79.8353 |
| 2737762 | -79.8306 |
| 84379317 | -79.8282 |
| 21737917 | -79.8083 |
| 14009261 | -79.7995 |
| 7693497 | -79.7907 |
| 10198899 | -79.7864 |
| 11818503 | -79.779 |
| 125493624 | -79.7662 |
| 57351860 | -79.7636 |
| 13696370 | -79.7508 |
| 46504433 | -79.7426 |
| 2749603 | -79.7411 |
| 235838 | -79.737 |
| 15102410 | -79.7188 |
| 334439 | -79.7158 |
| 57509304 | -79.7127 |
| 21437455 | -79.7054 |
| 71400828 | -79.7026 |
| 84375495 | -79.6933 |
| 20089828 | -79.6907 |
| 40468639 | -79.687 |
| 7106151 | -79.6834 |
| 354141 | -79.6815 |
| 71351029 | -79.675 |
| 53440807 | -79.6747 |
| 84379947 | -79.6608 |
| 20234790 | -79.6543 |
| 4531930 | -79.6496 |
| 56696870 | -79.6486 |
| 153369 | -79.6433 |
| 136270401 | -79.6349 |
| 2847808 | -79.6317 |
| 443788 | -79.6299 |
| 15118825 | -79.623 |
| 59054114 | -79.6155 |
| 13005615 | -79.6116 |
| 153074 | -79.604 |
| 71351426 | -79.5996 |
| 125745441 | -79.5958 |
| 2749882 | -79.5793 |
| 8219224 | -79.5676 |
| 13650419 | -79.5645 |
| 10947420 | -79.5547 |
| 12778437 | -79.5507 |
| 3358745 | -79.5419 |
| 189825 | -79.5364 |
| 12305200 | -79.5262 |
| 71402299 | -79.5246 |
| 11602229 | -79.5167 |
| 7481529 | -79.5137 |
| 16767195 | -79.5123 |
| 10825765 | -79.5113 |
| 3551245 | -79.4978 |
| 66521672 | -79.4781 |
| 626497 | -79.4778 |
| 7072969 | -79.4748 |
| 619375 | -79.474 |
| 124223113 | -79.472 |
| 68114119 | -79.4684 |
| 617280 | -79.4614 |
| 3089798 | -79.4512 |
| 98126733 | -79.4496 |
| 4355899 | -79.4476 |
| 333918 | -79.447 |
| 4988955 | -79.4411 |
| 12313954 | -79.4399 |
| 91664797 | -79.4371 |
| 18877558 | -79.4356 |
| 7345576 | -79.4279 |
| 18668027 | -79.424 |
| 56442 | -79.4175 |
| 15025286 | -79.4153 |
| 3848399 | -79.3995 |
| 438086 | -79.3972 |
| 128149 | -79.3887 |
| 160717 | -79.3743 |
| 363422 | -79.3665 |
| 14705051 | -79.3499 |
| 102507261 | -79.344 |
| 6454617 | -79.331 |
| 13945063 | -79.3297 |
| 12714658 | -79.3198 |
| 12318743 | -79.3189 |
| 19024166 | -79.3176 |
| 124062 | -79.3168 |
| 625612 | -79.3143 |
| 66489 | -79.2718 |
| 349039 | -79.2675 |
| 3083575 | -79.2661 |
| 629091 | -79.2611 |
| 15883490 | -79.2515 |
| 16205917 | -79.2488 |
| 15520942 | -79.2467 |
| 6454715 | -79.2409 |
| 12386616 | -79.2358 |
| 343025 | -79.226 |
| 238958 | -79.2231 |
| 17822020 | -79.2193 |
| 92538 | -79.2168 |
| 71358705 | -79.2164 |
| 428782 | -79.2127 |
| 623044 | -79.2104 |
| 106019 | -79.2059 |
| 10019885 | -79.2038 |
| 350382 | -79.2005 |
| 146050674 | -79.1997 |
| 7062292 | -79.195 |
| 6849333 | -79.1938 |
| 15519694 | -79.1901 |
| 3929905 | -79.1825 |
| 12342681 | -79.1778 |
| 13412789 | -79.1759 |
| 5315956 | -79.17 |
| 8032598 | -79.1686 |
| 124063 | -79.1665 |
| 131199 | -79.1663 |
| 15794174 | -79.1637 |
| 98115207 | -79.1534 |
| 5704920 | -79.1473 |
| 11031986 | -79.1351 |
| 88090596 | -79.1069 |
| 269955 | -79.0818 |
| 236908 | -79.0785 |
| 128452 | -79.0771 |
| 275493 | -79.0717 |
| 2870897 | -79.0672 |
| 3126032 | -79.0576 |
| 74147 | -79.0316 |
| 146825 | -79.0283 |
| 4003195 | -79.0234 |
| 91823808 | -79.021 |
| 3100885 | -79.0091 |
| 45490214 | -79.0052 |
| 66529 | -78.9941 |
| 125095 | -78.9928 |
| 423407 | -78.9928 |
| 7065338 | -78.9896 |
| 13897303 | -78.9857 |
| 56693987 | -78.9792 |
| 624969 | -78.9754 |
| 11771634 | -78.9706 |
| 2930198 | -78.969 |
| 3321714 | -78.9601 |
| 71344580 | -78.959 |
| 71349528 | -78.9517 |
| 44287782 | -78.9503 |
| 15742198 | -78.9464 |
| 16206119 | -78.9433 |
| 4455889 | -78.9415 |
| 95936 | -78.9406 |
| 189763 | -78.9366 |
| 12278638 | -78.9348 |
| 316854 | -78.922 |
| 13129200 | -78.9202 |
| 71365469 | -78.9163 |
| 7345499 | -78.9006 |
| 87489845 | -78.8997 |
| 91664814 | -78.8943 |
| 44287967 | -78.8935 |
| 71210144 | -78.8888 |
| 23986036 | -78.8869 |
| 5070780 | -78.8819 |
| 2917033 | -78.8677 |
| 22413850 | -78.8614 |
| 2857195 | -78.8597 |
| 18646 | -78.857 |
| 14320101 | -78.8524 |
| 232895 | -78.8514 |
| 71413104 | -78.8511 |
| 7063305 | -78.845 |
| 614116 | -78.8325 |
| 15269202 | -78.8307 |
| 2848043 | -78.8282 |
| 75530435 | -78.8212 |
| 624961 | -78.8166 |
| 13130520 | -78.8154 |
| 71367021 | -78.812 |
| 11768 | -78.8098 |
| 3155950 | -78.802 |
| 15651852 | -78.799 |
| 148265 | -78.7945 |
| 11833492 | -78.782 |
| 614108 | -78.7805 |
| 84376864 | -78.7773 |
| 482558 | -78.7737 |
| 11265324 | -78.7726 |
| 71402906 | -78.7684 |
| 3377640 | -78.7673 |
| 11161540 | -78.7667 |
| 3016789 | -78.7603 |
| 54708874 | -78.7546 |
| 44445519 | -78.75 |
| 168797 | -78.7477 |
| 7330510 | -78.7446 |
| 98539 | -78.7401 |
| 171106 | -78.7393 |
| 626498 | -78.7367 |
| 51136565 | -78.7345 |
| 71338074 | -78.7307 |
| 71349200 | -78.7303 |
| 71325692 | -78.728 |
| 94940646 | -78.7202 |
| 7324703 | -78.7129 |
| 2933273 | -78.7108 |
| 190004 | -78.7064 |
| 7230396 | -78.7046 |
| 16417531 | -78.6878 |
| 5318738 | -78.6805 |
| 13970503 | -78.6778 |
| 4135979 | -78.6677 |
| 202696 | -78.6672 |
| 384119 | -78.6613 |
| 2749879 | -78.6602 |
| 195147 | -78.6569 |
| 5323531 | -78.6566 |
| 620049 | -78.6516 |
| 71410867 | -78.647 |
| 426146 | -78.6366 |
| 71384878 | -78.6186 |
| 12497096 | -78.5847 |
| 13129226 | -78.5694 |
| 175017 | -78.5667 |
| 10912569 | -78.5644 |
| 10611710 | -78.5627 |
| 31533 | -78.5507 |
| 346739 | -78.539 |
| 14524514 | -78.5331 |
| 3906062 | -78.5212 |
| 12497098 | -78.5166 |
| 66521666 | -78.5136 |
| 7060273 | -78.5107 |
| 624912 | -78.4873 |
| 7106152 | -78.4869 |
| 13182760 | -78.4767 |
| 620549 | -78.475 |
| 10152 | -78.4722 |
| 627335 | -78.4677 |
| 21419215 | -78.4455 |
| 274333 | -78.4426 |
| 80103 | -78.4361 |
| 131861694 | -78.4359 |
| 106143 | -78.4337 |
| 85919892 | -78.4274 |
| 22643677 | -78.4269 |
| 354138 | -78.4266 |
| 313347 | -78.4259 |
| 6683 | -78.424 |
| 41930 | -78.424 |
| 12392754 | -78.4215 |
| 132521 | -78.4125 |
| 18927945 | -78.4109 |
| 12804073 | -78.3926 |
| 318666 | -78.3873 |
| 124869 | -78.3806 |
| 12963549 | -78.3755 |
| 20528 | -78.3753 |
| 153102 | -78.3644 |
| 350272 | -78.3632 |
| 438245 | -78.3603 |
| 439019 | -78.3603 |
| 53249899 | -78.3451 |
| 161310 | -78.3428 |
| 316855 | -78.3406 |
| 2748951 | -78.3393 |
| 11779716 | -78.3364 |
| 4173710 | -78.336 |
| 29940170 | -78.3347 |
| 3952843 | -78.3327 |
| 347430 | -78.3286 |
| 71323793 | -78.3088 |
| 3501064 | -78.3002 |
| 10902125 | -78.2993 |
| 12262011 | -78.2957 |
| 103214 | -78.2939 |
| 3126049 | -78.2914 |
| 85977017 | -78.2884 |
| 10169 | -78.2734 |
| 12546955 | -78.2684 |
| 85932629 | -78.262 |
| 71316319 | -78.2569 |
| 3010923 | -78.2411 |
| 5035038 | -78.2324 |
| 12342682 | -78.2313 |
| 73894533 | -78.2307 |
| 160474 | -78.2242 |
| 111408 | -78.2209 |
| 95861 | -78.2194 |
| 14419219 | -78.2111 |
| 3912165 | -78.2097 |
| 5761855 | -78.2085 |
| 6710629 | -78.2034 |
| 10730442 | -78.2 |
| 81473 | -78.1859 |
| 5004 | -78.1813 |
| 458184 | -78.1796 |
| 92278763 | -78.1686 |
| 91665183 | -78.1622 |
| 640886 | -78.1488 |
| 69440 | -78.1476 |
| 2888130 | -78.1428 |
| 71327521 | -78.1092 |
| 12329386 | -78.1023 |
| 151887 | -78.0974 |
| 71386630 | -78.0793 |
| 3833577 | -78.0759 |
| 3220 | -78.0684 |
| 112840 | -78.0551 |
| 191290 | -78.0465 |
| 22026216 | -78.0433 |
| 92258675 | -78.0398 |
| 2752529 | -78.0358 |
| 12220564 | -78.0343 |
| 42648414 | -78.0235 |
| 24787792 | -78.0151 |
| 3545630 | -78.012 |
| 40553282 | -78.0009 |
| 20568309 | -78.0003 |
| 118430 | -77.998 |
| 236911 | -77.9941 |
| 2857011 | -77.9938 |
| 95931189 | -77.9936 |
| 7063304 | -77.9828 |
| 4374653 | -77.9604 |
| 15204558 | -77.9542 |
| 146675964 | -77.9535 |
| 91665190 | -77.9301 |
| 626486 | -77.9269 |
| 3100888 | -77.9187 |
| 15559331 | -77.9185 |
| 11836860 | -77.9183 |
| 94938899 | -77.9173 |
| 12304766 | -77.9088 |
| 22643725 | -77.8879 |
| 18877559 | -77.874 |
| 71372876 | -77.8632 |
| 5704988 | -77.8608 |
| 235827 | -77.8538 |
| 71403676 | -77.8406 |
| 71399352 | -77.8369 |
| 12260949 | -77.832 |
| 12454279 | -77.818 |
| 124464351 | -77.8156 |
| 13051574 | -77.8112 |
| 13245081 | -77.8067 |
| 23677362 | -77.8032 |
| 21953556 | -77.7966 |
| 442767 | -77.7954 |
| 2847380 | -77.7936 |
| 17999326 | -77.7742 |
| 12756415 | -77.7681 |
| 9819207 | -77.7645 |
| 5146503 | -77.7629 |
| 16205214 | -77.7448 |
| 1320 | -77.7333 |
| 626208 | -77.7292 |
| 4388935 | -77.7231 |
| 428781 | -77.7172 |
| 147143 | -77.7161 |
| 57368489 | -77.7098 |
| 23423990 | -77.7087 |
| 90481772 | -77.6969 |
| 3086089 | -77.6931 |
| 27000054 | -77.6828 |
| 92283377 | -77.6809 |
| 58434465 | -77.6706 |
| 125464518 | -77.6655 |
| 620298 | -77.6646 |
| 151618 | -77.6574 |
| 5325701 | -77.6555 |
| 13291774 | -77.6398 |
| 44251290 | -77.6332 |
| 85686311 | -77.6326 |
| 100939938 | -77.6292 |
| 10661597 | -77.629 |
| 85686314 | -77.6179 |
| 2835176 | -77.6165 |
| 271593 | -77.6152 |
| 3886438 | -77.614 |
| 188082 | -77.6003 |
| 2749858 | -77.5987 |
| 44715573 | -77.5973 |
| 21794841 | -77.5873 |
| 12347528 | -77.5771 |
| 71386138 | -77.5754 |
| 67212 | -77.5739 |
| 20362533 | -77.5735 |
| 71353500 | -77.5735 |
| 5325700 | -77.5713 |
| 435950 | -77.5685 |
| 342664 | -77.5592 |
| 71319045 | -77.5565 |
| 4530845 | -77.5541 |
| 71319024 | -77.5524 |
| 434660 | -77.5485 |
| 13082567 | -77.5434 |
| 16051915 | -77.5431 |
| 458830 | -77.5397 |
| 21575659 | -77.5374 |
| 65739 | -77.532 |
| 16765149 | -77.5316 |
| 9992552 | -77.5306 |
| 4125339 | -77.5217 |
| 148954 | -77.5155 |
| 12322287 | -77.5143 |
| 436367 | -77.5088 |
| 3100886 | -77.5023 |
| 23670790 | -77.4934 |
| 11044279 | -77.4894 |
| 5019413 | -77.4878 |
| 3725402 | -77.4869 |
| 15778411 | -77.486 |
| 4143813 | -77.4818 |
| 14251911 | -77.4803 |
| 133561627 | -77.4771 |
| 554081 | -77.477 |
| 44044 | -77.4666 |
| 437847 | -77.4584 |
| 2916983 | -77.4569 |
| 71425079 | -77.4564 |
| 56692412 | -77.4513 |
| 2849150 | -77.4494 |
| 4996383 | -77.4454 |
| 59942142 | -77.4241 |
| 3019962 | -77.4231 |
| 4395282 | -77.4216 |
| 44043 | -77.4061 |
| 361511 | -77.4059 |
| 2879318 | -77.4016 |
| 67392128 | -77.3793 |
| 147356 | -77.3775 |
| 61719 | -77.3757 |
| 412282 | -77.3726 |
| 71406968 | -77.3724 |
| 91886385 | -77.3701 |
| 5182247 | -77.3646 |
| 71386136 | -77.3609 |
| 15650021 | -77.353 |
| 85990596 | -77.3485 |
| 425993 | -77.3462 |
| 12780391 | -77.3329 |
| 76415 | -77.3168 |
| 71365213 | -77.3072 |
| 3664123 | -77.301 |
| 12634029 | -77.2699 |
| 46781449 | -77.2662 |
| 419290 | -77.2638 |
| 21773239 | -77.2522 |
| 20291130 | -77.2316 |
| 407615 | -77.231 |
| 160475 | -77.2295 |
| 54720345 | -77.2156 |
| 3770390 | -77.2109 |
| 80059 | -77.1911 |
| 184381 | -77.1822 |
| 343740 | -77.1817 |
| 92246578 | -77.1766 |
| 7209385 | -77.1734 |
| 131899 | -77.1729 |
| 92278714 | -77.1687 |
| 13455967 | -77.167 |
| 21497699 | -77.1406 |
| 51058474 | -77.1396 |
| 160483 | -77.1395 |
| 97560 | -77.1188 |
| 125495998 | -77.1159 |
| 19987802 | -77.1137 |
| 12307112 | -77.1133 |
| 458225 | -77.1131 |
| 317023 | -77.095 |
| 112865 | -77.093 |
| 15200919 | -77.0885 |
| 5338491 | -77.0724 |
| 10912983 | -77.07 |
| 5704921 | -77.0652 |
| 354142 | -77.0631 |
| 24972908 | -77.0551 |
| 15138599 | -77.053 |
| 71324957 | -77.05 |
| 419323 | -77.0499 |
| 112237 | -77.0337 |
| 13129224 | -77.0334 |
| 15998304 | -77.023 |
| 82200940 | -77.0217 |
| 71379714 | -77.0203 |
| 273071 | -77.019 |
| 41933 | -77.0154 |
| 12730689 | -77.0143 |
| 7209528 | -77.0117 |
| 41935 | -77.0075 |
| 4205046 | -76.9983 |
| 2890949 | -76.9966 |
| 71386133 | -76.9894 |
| 71332328 | -76.9857 |
| 19729 | -76.9836 |
| 236865 | -76.9829 |
| 2760431 | -76.951 |
| 425893 | -76.9432 |
| 25021 | -76.9431 |
| 23271 | -76.9397 |
| 42586888 | -76.9342 |
| 21413445 | -76.9326 |
| 68235121 | -76.925 |
| 301142 | -76.9204 |
| 12506460 | -76.9171 |
| 4412951 | -76.9168 |
| 442759 | -76.9146 |
| 271597 | -76.9118 |
| 13360158 | -76.8989 |
| 294389 | -76.8901 |
| 443810 | -76.8871 |
| 23032 | -76.8818 |
| 12825803 | -76.8774 |
| 7694432 | -76.8758 |
| 13659965 | -76.8682 |
| 12220563 | -76.8581 |
| 10149918 | -76.8538 |
| 3695121 | -76.8505 |
| 354139 | -76.84 |
| 344329 | -76.8398 |
| 20642869 | -76.8334 |
| 3025796 | -76.8242 |
| 131862131 | -76.8226 |
| 71394847 | -76.8204 |
| 84376862 | -76.8188 |
| 53440717 | -76.7982 |
| 14048786 | -76.7947 |
| 348101 | -76.7867 |
| 3440007 | -76.785 |
| 277276 | -76.7824 |
| 3085033 | -76.781 |
| 12659857 | -76.7781 |
| 15993164 | -76.777 |
| 5184418 | -76.7759 |
| 7721310 | -76.7737 |
| 91664808 | -76.7717 |
| 11067354 | -76.7713 |
| 10151 | -76.7654 |
| 12672702 | -76.7581 |
| 56443 | -76.7289 |
| 116939 | -76.7235 |
| 71386132 | -76.7228 |
| 12007654 | -76.7223 |
| 71441837 | -76.7116 |
| 71359083 | -76.7011 |
| 639112 | -76.6868 |
| 2847322 | -76.6823 |
| 179447 | -76.6619 |
| 10150 | -76.6616 |
| 146027 | -76.6557 |
| 15579026 | -76.6497 |
| 4664282 | -76.6315 |
| 15993104 | -76.6306 |
| 10208 | -76.6195 |
| 12628832 | -76.6139 |
| 554074 | -76.5783 |
| 66959208 | -76.5715 |
| 213193 | -76.5677 |
| 125585 | -76.5652 |
| 314228 | -76.5546 |
| 9541459 | -76.554 |
| 5171130 | -76.5464 |
| 71406207 | -76.5461 |
| 21419216 | -76.5442 |
| 5062045 | -76.5411 |
| 53745780 | -76.5334 |
| 16640709 | -76.5308 |
| 10400914 | -76.5214 |
| 16769826 | -76.5133 |
| 7980493 | -76.5075 |
| 71330484 | -76.4873 |
| 15743472 | -76.4852 |
| 12307090 | -76.4684 |
| 12302772 | -76.467 |
| 350510 | -76.4661 |
| 11065672 | -76.4661 |
| 8032526 | -76.4624 |
| 53424595 | -76.4601 |
| 7122776 | -76.4599 |
| 13129225 | -76.4596 |
| 5318136 | -76.45 |
| 350512 | -76.4493 |
| 14320097 | -76.4321 |
| 6682 | -76.4303 |
| 15579048 | -76.4273 |
| 80761 | -76.4163 |
| 4439825 | -76.4094 |
| 13675106 | -76.4058 |
| 626944 | -76.4032 |
| 11230545 | -76.3959 |
| 363263 | -76.3742 |
| 135493748 | -76.3713 |
| 2833794 | -76.3629 |
| 12322346 | -76.3626 |
| 20095224 | -76.361 |
| 100242 | -76.3582 |
| 17282406 | -76.3519 |
| 15259337 | -76.3472 |
| 11196859 | -76.3447 |
| 131843860 | -76.3398 |
| 11107397 | -76.3345 |
| 75433393 | -76.3342 |
| 71357852 | -76.3331 |
| 131851433 | -76.3233 |
| 4624554 | -76.2994 |
| 21318740 | -76.2984 |
| 12306672 | -76.2921 |
| 11196140 | -76.2742 |
| 22643716 | -76.2647 |
| 10514946 | -76.2564 |
| 12360345 | -76.2486 |
| 13130307 | -76.2448 |
| 18593 | -76.2246 |
| 42647721 | -76.2194 |
| 429241 | -76.2147 |
| 17253 | -76.2078 |
| 13930872 | -76.2048 |
| 17189 | -76.1789 |
| 617886 | -76.1713 |
| 12278639 | -76.1709 |
| 66480 | -76.1667 |
| 4594253 | -76.1643 |
| 426675 | -76.1607 |
| 71386139 | -76.155 |
| 162276 | -76.1363 |
| 91903 | -76.1291 |
| 236872 | -76.1226 |
| 41932 | -76.1084 |
| 2849112 | -76.0884 |
| 617969 | -76.0632 |
| 620286 | -76.0514 |
| 6705 | -76.0497 |
| 12861652 | -76.0416 |
| 4561382 | -76.0301 |
| 71343455 | -76.0285 |
| 23260675 | -76.0258 |
| 410191 | -76.0185 |
| 12435249 | -76.0178 |
| 50601 | -76.0036 |
| 458174 | -76.0006 |
| 15946825 | -75.9862 |
| 235564 | -75.9822 |
| 624558 | -75.9687 |
| 71386631 | -75.9677 |
| 71329043 | -75.9588 |
| 3301305 | -75.9586 |
| 12276809 | -75.9554 |
| 21120631 | -75.9483 |
| 155896787 | -75.941 |
| 66508 | -75.9332 |
| 631456 | -75.9327 |
| 301143 | -75.9288 |
| 5319478 | -75.9202 |
| 83425 | -75.9194 |
| 2887738 | -75.9057 |
| 336581 | -75.8853 |
| 7693389 | -75.8766 |
| 76071 | -75.8746 |
| 46911017 | -75.8713 |
| 6451286 | -75.8709 |
| 2948 | -75.8687 |
| 20324363 | -75.8653 |
| 96191 | -75.8631 |
| 12164257 | -75.848 |
| 3807128 | -75.8479 |
| 10148 | -75.8477 |
| 12233923 | -75.8421 |
| 122458 | -75.8216 |
| 21520019 | -75.8191 |
| 44538 | -75.8149 |
| 2749308 | -75.8069 |
| 12307119 | -75.7974 |
| 11254074 | -75.7968 |
| 8032574 | -75.7825 |
| 11253808 | -75.7706 |
| 314049 | -75.7699 |
| 13945060 | -75.7665 |
| 40468612 | -75.7554 |
| 4156291 | -75.755 |
| 66510 | -75.7421 |
| 3915367 | -75.7374 |
| 3256725 | -75.7297 |
| 12307117 | -75.7287 |
| 304457 | -75.7246 |
| 12642744 | -75.7198 |
| 72910 | -75.7195 |
| 19429813 | -75.7137 |
| 637241 | -75.7087 |
| 103062 | -75.6975 |
| 2749865 | -75.697 |
| 71324956 | -75.6888 |
| 12435251 | -75.6846 |
| 3993156 | -75.6768 |
| 13050218 | -75.6736 |
| 12567021 | -75.6711 |
| 622060 | -75.6697 |
| 66509 | -75.6452 |
| 100574 | -75.6352 |
| 23194572 | -75.6313 |
| 13257589 | -75.6251 |
| 40502 | -75.6153 |
| 184386 | -75.6013 |
| 13455968 | -75.5976 |
| 71397610 | -75.5935 |
| 126957664 | -75.58 |
| 517266 | -75.5793 |
| 44784496 | -75.5712 |
| 458177 | -75.5684 |
| 638851 | -75.5627 |
| 118740 | -75.5507 |
| 42648293 | -75.5255 |
| 99300 | -75.5036 |
| 220107 | -75.4884 |
| 15994522 | -75.4846 |
| 458173 | -75.4583 |
| 10087355 | -75.4561 |
| 19154573 | -75.45 |
| 7980495 | -75.4368 |
| 3905449 | -75.4296 |
| 12325858 | -75.4089 |
| 4534766 | -75.3999 |
| 12307120 | -75.3838 |
| 12804072 | -75.3796 |
| 5319467 | -75.3786 |
| 5325698 | -75.371 |
| 12744373 | -75.3691 |
| 67194 | -75.3681 |
| 4161864 | -75.3609 |
| 85788115 | -75.3565 |
| 7730236 | -75.3541 |
| 78237 | -75.3496 |
| 67192 | -75.3435 |
| 3547804 | -75.3431 |
| 12659855 | -75.3419 |
| 4594254 | -75.3357 |
| 2840324 | -75.3346 |
| 11738774 | -75.3211 |
| 10879636 | -75.3023 |
| 236874 | -75.2934 |
| 23511067 | -75.2889 |
| 156023563 | -75.2761 |
| 2748950 | -75.2621 |
| 11300144 | -75.2535 |
| 314232 | -75.2487 |
| 437525 | -75.2351 |
| 438001 | -75.2328 |
| 32208 | -75.23 |
| 182384 | -75.2272 |
| 57675852 | -75.2208 |
| 71347226 | -75.2155 |
| 5244434 | -75.2121 |
| 12332070 | -75.2029 |
| 5485989 | -75.2028 |
| 2916838 | -75.2025 |
| 71356129 | -75.1979 |
| 21847321 | -75.1952 |
| 619891 | -75.1899 |
| 22849 | -75.1793 |
| 350508 | -75.1792 |
| 160817 | -75.1788 |
| 21403139 | -75.1642 |
| 4314659 | -75.1456 |
| 12334638 | -75.1353 |
| 12436930 | -75.134 |
| 20095231 | -75.1328 |
| 13303585 | -75.1314 |
| 11300870 | -75.1311 |
| 11482554 | -75.131 |
| 12262032 | -75.127 |
| 10268224 | -75.1089 |
| 71357686 | -75.1002 |
| 3126043 | -75.093 |
| 3919311 | -75.0867 |
| 108921 | -75.0752 |
| 94037499 | -75.0689 |
| 12744370 | -75.0673 |
| 22952 | -75.0663 |
| 71370767 | -75.0657 |
| 313900 | -75.0563 |
| 53431632 | -75.0496 |
| 11107822 | -75.0446 |
| 6451440 | -75.0337 |
| 236907 | -75.0262 |
| 281333 | -75.024 |
| 3544983 | -75.0108 |
| 20194334 | -74.9813 |
| 66482 | -74.9797 |
| 11391150 | -74.9761 |
| 11522009 | -74.9632 |
| 14982738 | -74.9492 |
| 5704844 | -74.9326 |
| 71386130 | -74.9304 |
| 9883616 | -74.9211 |
| 12459209 | -74.9156 |
| 3106277 | -74.913 |
| 2749315 | -74.9073 |
| 4112659 | -74.8912 |
| 80309 | -74.8876 |
| 351877 | -74.8841 |
| 76295 | -74.8804 |
| 15557501 | -74.8772 |
| 9581277 | -74.8631 |
| 12644531 | -74.8573 |
| 71337652 | -74.8566 |
| 16205060 | -74.851 |
| 104188 | -74.8494 |
| 236873 | -74.8488 |
| 71372099 | -74.8437 |
| 40019 | -74.8412 |
| 12314177 | -74.8392 |
| 54444107 | -74.8304 |
| 14160 | -74.8301 |
| 12635763 | -74.8222 |
| 10683232 | -74.8185 |
| 71399353 | -74.8163 |
| 12812113 | -74.8159 |
| 71324955 | -74.8119 |
| 91738908 | -74.8108 |
| 83633 | -74.8018 |
| 182449 | -74.7984 |
| 2837523 | -74.7975 |
| 3910963 | -74.7957 |
| 6704 | -74.7861 |
| 15561151 | -74.785 |
| 76695 | -74.771 |
| 459274 | -74.7689 |
| 412448 | -74.7649 |
| 7694433 | -74.7631 |
| 617285 | -74.7616 |
| 617701 | -74.7456 |
| 71399351 | -74.7369 |
| 10778215 | -74.7258 |
| 12891049 | -74.7244 |
| 236894 | -74.722 |
| 196978 | -74.7187 |
| 13412788 | -74.7112 |
| 23325142 | -74.7089 |
| 117969 | -74.7061 |
| 3085651 | -74.7009 |
| 53240291 | -74.6918 |
| 68445 | -74.6751 |
| 23267239 | -74.6609 |
| 2834683 | -74.6593 |
| 363560 | -74.656 |
| 20536776 | -74.6446 |
| 99641155 | -74.6246 |
| 3090608 | -74.6071 |
| 13668577 | -74.5884 |
| 84896 | -74.5847 |
| 131861813 | -74.5773 |
| 69166000 | -74.5767 |
| 186840 | -74.5746 |
| 10537883 | -74.5668 |
| 9817583 | -74.5648 |
| 59253883 | -74.5621 |
| 12384202 | -74.5581 |
| 150722 | -74.5512 |
| 458833 | -74.5448 |
| 71368150 | -74.5428 |
| 12678918 | -74.5182 |
| 19833424 | -74.5151 |
| 81989 | -74.5145 |
| 59253871 | -74.5029 |
| 12317190 | -74.4741 |
| 13781990 | -74.4691 |
| 3527267 | -74.4506 |
| 71394212 | -74.4301 |
| 4130196 | -74.4263 |
| 12400976 | -74.4135 |
| 11796 | -74.4009 |
| 145711456 | -74.3874 |
| 13197044 | -74.3845 |
| 80827 | -74.3834 |
| 350537 | -74.3784 |
| 459271 | -74.3784 |
| 11833885 | -74.3751 |
| 20240718 | -74.3546 |
| 3440575 | -74.3444 |
| 14199976 | -74.3403 |
| 6454445 | -74.3343 |
| 91904 | -74.3309 |
| 69916862 | -74.3283 |
| 95005217 | -74.3178 |
| 3085855 | -74.3046 |
| 22708978 | -74.3028 |
| 66481 | -74.301 |
| 122909 | -74.2992 |
| 4140533 | -74.2912 |
| 57351859 | -74.2672 |
| 22643718 | -74.2489 |
| 71365214 | -74.2224 |
| 5152711 | -74.2198 |
| 6454985 | -74.2164 |
| 622701 | -74.1881 |
| 24181320 | -74.1777 |
| 20502346 | -74.1773 |
| 13056536 | -74.1625 |
| 71386134 | -74.1558 |
| 10891675 | -74.1265 |
| 31421 | -74.1192 |
| 12224293 | -74.116 |
| 15579021 | -74.1114 |
| 57351858 | -74.0989 |
| 3141101 | -74.0959 |
| 108957 | -74.0897 |
| 12641085 | -74.084 |
| 8322 | -74.0825 |
| 71368906 | -74.078 |
| 14323661 | -74.0697 |
| 3955344 | -74.0352 |
| 152630 | -74.0224 |
| 76338 | -74.0105 |
| 24186919 | -74.0093 |
| 82030669 | -74.0049 |
| 4408246 | -73.9991 |
| 22321051 | -73.9894 |
| 5247620 | -73.9762 |
| 71379711 | -73.9661 |
| 7062418 | -73.9513 |
| 91665189 | -73.9489 |
| 40568053 | -73.9417 |
| 71365594 | -73.9267 |
| 3678091 | -73.8811 |
| 15570414 | -73.8705 |
| 623876 | -73.8544 |
| 71413590 | -73.8287 |
| 71373383 | -73.7767 |
| 8510 | -73.7719 |
| 1382 | -73.7695 |
| 3020421 | -73.7655 |
| 2749312 | -73.7606 |
| 12531986 | -73.7352 |
| 51138612 | -73.7287 |
| 10610487 | -73.7279 |
| 145710580 | -73.7275 |
| 16205286 | -73.7219 |
| 16205119 | -73.711 |
| 266054 | -73.7008 |
| 4421615 | -73.6746 |
| 71380657 | -73.673 |
| 8534 | -73.6488 |
| 626024 | -73.6451 |
| 2917272 | -73.6425 |
| 3534338 | -73.6361 |
| 15716481 | -73.6024 |
| 71367052 | -73.5985 |
| 12628831 | -73.5963 |
| 13439296 | -73.5788 |
| 236904 | -73.5765 |
| 12963547 | -73.5682 |
| 459343 | -73.5618 |
| 626560 | -73.5568 |
| 3554357 | -73.535 |
| 13621081 | -73.5327 |
| 11798 | -73.5292 |
| 17384162 | -73.5229 |
| 85773419 | -73.5174 |
| 20521451 | -73.5048 |
| 3090511 | -73.5047 |
| 236909 | -73.4867 |
| 71299271 | -73.4827 |
| 344311 | -73.472 |
| 11536666 | -73.4647 |
| 6714 | -73.4643 |
| 3438280 | -73.4619 |
| 12531982 | -73.4563 |
| 2833772 | -73.4374 |
| 236869 | -73.433 |
| 434596 | -73.4328 |
| 53435541 | -73.4319 |
| 616789 | -73.4286 |
| 83344 | -73.4222 |
| 98164465 | -73.4183 |
| 131851613 | -73.3973 |
| 4657174 | -73.3877 |
| 2840206 | -73.3871 |
| 71042 | -73.3761 |
| 344452 | -73.3674 |
| 80953 | -73.3615 |
| 3881848 | -73.3497 |
| 5182448 | -73.3081 |
| 13906820 | -73.3062 |
| 235820 | -73.3044 |
| 155237 | -73.3 |
| 12772513 | -73.2976 |
| 258553 | -73.2887 |
| 346879 | -73.2803 |
| 4634231 | -73.2785 |
| 2840432 | -73.278 |
| 5464551 | -73.2771 |
| 151616 | -73.2767 |
| 179389 | -73.2716 |
| 2764161 | -73.2488 |
| 129646071 | -73.2346 |
| 4153129 | -73.2344 |
| 18551879 | -73.2219 |
| 20034870 | -73.1851 |
| 15556316 | -73.1819 |
| 4311527 | -73.1668 |
| 164982 | -73.1604 |
| 33790 | -73.1561 |
| 235575 | -73.1559 |
| 6776 | -73.1476 |
| 14886011 | -73.1457 |
| 56694105 | -73.133 |
| 8557 | -73.0952 |
| 12262031 | -73.0591 |
| 3100909 | -73.0455 |
| 2749876 | -73.0229 |
| 80769 | -73.0057 |
| 104647 | -73.0043 |
| 7060303 | -72.9846 |
| 2749855 | -72.9802 |
| 13944287 | -72.9769 |
| 2847694 | -72.9539 |
| 236900 | -72.9452 |
| 10490094 | -72.9313 |
| 71370710 | -72.9303 |
| 131859373 | -72.9164 |
| 623980 | -72.9131 |
| 47560 | -72.9073 |
| 100818169 | -72.8848 |
| 99637906 | -72.8712 |
| 620996 | -72.8693 |
| 13593656 | -72.8683 |
| 10889963 | -72.8678 |
| 10911747 | -72.8497 |
| 124928536 | -72.8294 |
| 69856147 | -72.8239 |
| 14274944 | -72.8012 |
| 12678916 | -72.7972 |
| 220594 | -72.7858 |
| 91665184 | -72.7762 |
| 13245082 | -72.7754 |
| 67210 | -72.7751 |
| 12307094 | -72.7727 |
| 5316800 | -72.7682 |
| 53719213 | -72.7629 |
| 6702 | -72.7597 |
| 3090916 | -72.7345 |
| 2749908 | -72.7313 |
| 3150757 | -72.7214 |
| 131862030 | -72.7166 |
| 71316217 | -72.7086 |
| 20034887 | -72.7043 |
| 131858361 | -72.6869 |
| 57360689 | -72.6777 |
| 12631756 | -72.6745 |
| 131861234 | -72.67 |
| 620609 | -72.6667 |
| 236896 | -72.6631 |
| 71357987 | -72.6486 |
| 7377958 | -72.6449 |
| 412444 | -72.6421 |
| 19887666 | -72.6406 |
| 2950 | -72.6346 |
| 131860067 | -72.622 |
| 20521443 | -72.6161 |
| 6293 | -72.6159 |
| 78544 | -72.6133 |
| 433774 | -72.6038 |
| 458954 | -72.6025 |
| 13796464 | -72.5977 |
| 3106281 | -72.5971 |
| 8321 | -72.5949 |
| 25528 | -72.5859 |
| 53438721 | -72.5727 |
| 42647720 | -72.5666 |
| 12336760 | -72.5468 |
| 2840104 | -72.5392 |
| 85466 | -72.5336 |
| 4672069 | -72.5265 |
| 412063 | -72.5256 |
| 3563882 | -72.5216 |
| 22643753 | -72.5077 |
| 75590 | -72.5069 |
| 131859372 | -72.5042 |
| 2840268 | -72.5028 |
| 41931 | -72.4906 |
| 67029 | -72.485 |
| 7345500 | -72.4786 |
| 6711 | -72.4399 |
| 31418 | -72.438 |
| 3090513 | -72.4362 |
| 6688 | -72.4348 |
| 641675 | -72.4164 |
| 8512 | -72.4041 |
| 12336759 | -72.4036 |
| 2847772 | -72.3952 |
| 6772 | -72.3916 |
| 344316 | -72.3757 |
| 104142 | -72.3537 |
| 73072 | -72.3536 |
| 3617264 | -72.3397 |
| 8328 | -72.3345 |
| 412445 | -72.3289 |
| 281335 | -72.315 |
| 333260 | -72.3136 |
| 350483 | -72.312 |
| 12333787 | -72.2998 |
| 101691020 | -72.2484 |
| 5192314 | -72.2205 |
| 271596 | -72.212 |
| 4639769 | -72.2109 |
| 88094 | -72.1984 |
| 6686 | -72.1902 |
| 363003 | -72.1884 |
| 6454980 | -72.1759 |
| 74473 | -72.1756 |
| 24185353 | -72.1723 |
| 2939293 | -72.1712 |
| 3090510 | -72.149 |
| 76760 | -72.1461 |
| 3146258 | -72.1423 |
| 7154473 | -72.1313 |
| 10730441 | -72.1256 |
| 23707848 | -72.1162 |
| 344301 | -72.1153 |
| 20205649 | -72.1102 |
| 627884 | -72.1068 |
| 124925681 | -72.1047 |
| 50998459 | -72.0943 |
| 6708 | -72.0768 |
| 235830 | -72.0663 |
| 6707 | -72.055 |
| 314639 | -72.054 |
| 100642 | -72.0448 |
| 88212 | -72.0374 |
| 3478173 | -72.0277 |
| 12444 | -72.0249 |
| 3654897 | -72.0243 |
| 4202956 | -72.0226 |
| 22806 | -72.0089 |
| 622037 | -72.0074 |
| 16205213 | -72.0016 |
| 12079598 | -71.9923 |
| 53422966 | -71.9823 |
| 236897 | -71.9756 |
| 221013 | -71.953 |
| 96470094 | -71.9418 |
| 12333786 | -71.9397 |
| 8553 | -71.9293 |
| 5020936 | -71.9223 |
| 18510005 | -71.9156 |
| 3090899 | -71.9115 |
| 31419 | -71.91 |
| 6773 | -71.8976 |
| 15652 | -71.8796 |
| 13182761 | -71.8752 |
| 627326 | -71.8738 |
| 17750895 | -71.8714 |
| 17884 | -71.8705 |
| 13725949 | -71.866 |
| 235821 | -71.8536 |
| 19897935 | -71.8484 |
| 124926973 | -71.8433 |
| 24185030 | -71.8386 |
| 5044326 | -71.8186 |
| 3116509 | -71.8071 |
| 6709 | -71.787 |
| 71399366 | -71.7804 |
| 56605805 | -71.7616 |
| 12260948 | -71.7574 |
| 616802 | -71.7308 |
| 3378440 | -71.7278 |
| 67012 | -71.7071 |
| 80636 | -71.6974 |
| 235565 | -71.6801 |
| 225704 | -71.6779 |
| 21413437 | -71.6683 |
| 71327485 | -71.661 |
| 317828 | -71.6606 |
| 20467724 | -71.6583 |
| 220725 | -71.6314 |
| 459273 | -71.5837 |
| 71386137 | -71.5834 |
| 5007344 | -71.5773 |
| 6701 | -71.5535 |
| 71414849 | -71.5507 |
| 14919032 | -71.5482 |
| 8509 | -71.5446 |
| 12079599 | -71.5425 |
| 246666 | -71.5371 |
| 8320 | -71.5324 |
| 13955281 | -71.501 |
| 236910 | -71.4525 |
| 4291009 | -71.4307 |
| 16927 | -71.4286 |
| 20188060 | -71.4238 |
| 22058815 | -71.4231 |
| 19422402 | -71.3915 |
| 71414848 | -71.383 |
| 70522837 | -71.3778 |
| 10890992 | -71.3633 |
| 70615403 | -71.354 |
| 223669 | -71.3525 |
| 33774 | -71.3475 |
| 5966445 | -71.3459 |
| 222568 | -71.338 |
| 3490678 | -71.3374 |
| 70379 | -71.3349 |
| 71386131 | -71.3185 |
| 4532120 | -71.311 |
| 42596 | -71.3021 |
| 2749871 | -71.2731 |
| 71400931 | -71.2523 |
| 6706 | -71.2324 |
| 2749909 | -71.2195 |
| 126038 | -71.2133 |
| 4367845 | -71.2072 |
| 56694103 | -71.2037 |
| 12344381 | -71.1859 |
| 3383102 | -71.1695 |
| 7731174 | -71.1342 |
| 56694104 | -71.1027 |
| 8323 | -71.0937 |
| 16205948 | -71.0889 |
| 13736896 | -71.0466 |
| 67351 | -71.0413 |
| 18362412 | -71.0361 |
| 433775 | -70.9719 |
| 68436 | -70.9668 |
| 13227199 | -70.9612 |
| 236895 | -70.9174 |
| 70614910 | -70.9156 |
| 71357957 | -70.9084 |
| 13897903 | -70.9017 |
| 53422417 | -70.9005 |
| 21520715 | -70.8772 |
| 363753 | -70.869 |
| 3863551 | -70.8385 |
| 6713 | -70.8021 |
| 4107356 | -70.7724 |
| 12631754 | -70.7593 |
| 12468599 | -70.7503 |
| 293266 | -70.7402 |
| 4208919 | -70.7278 |
| 66477 | -70.7262 |
| 73698 | -70.701 |
| 619554 | -70.6972 |
| 2749910 | -70.6703 |
| 71386135 | -70.6692 |
| 236863 | -70.6338 |
| 95859 | -70.6298 |
| 344130 | -70.6232 |
| 7184048 | -70.5707 |
| 17885 | -70.5647 |
| 15716476 | -70.5626 |
| 4179424 | -70.5555 |
| 7731179 | -70.5555 |
| 5359678 | -70.5315 |
| 357626 | -70.5234 |
| 3932515 | -70.5061 |
| 19773428 | -70.498 |
| 2749907 | -70.4922 |
| 131851614 | -70.4563 |
| 177551 | -70.4507 |
| 13011088 | -70.4433 |
| 66602384 | -70.3979 |
| 66602383 | -70.3576 |
| 21292553 | -70.3046 |
| 4637536 | -70.3038 |
| 23465320 | -70.3015 |
| 4306675 | -70.2814 |
| 14848086 | -70.2709 |
| 2818125 | -70.223 |
| 31426 | -70.2025 |
| 20456766 | -70.1941 |
| 2833793 | -70.147 |
| 87240251 | -70.1385 |
| 53421626 | -70.1373 |
| 4119984 | -70.1307 |
| 344300 | -70.1225 |
| 2757724 | -70.1193 |
| 24181284 | -70.102 |
| 87435928 | -70.1007 |
| 412064 | -70.0722 |
| 31420 | -70.0555 |
| 5192315 | -70.0473 |
| 12812111 | -70.0261 |
| 66532 | -70.013 |
| 355492 | -70.0084 |
| 54352181 | -69.9849 |
| 7345225 | -69.9714 |
| 12387998 | -69.9674 |
| 23702632 | -69.9633 |
| 12262010 | -69.96 |
| 6771 | -69.9485 |
| 88921653 | -69.9275 |
| 5361513 | -69.9227 |
| 8341 | -69.9043 |
| 12860190 | -69.8817 |
| 82420 | -69.8768 |
| 109808 | -69.8542 |
| 131862135 | -69.8483 |
| 21403137 | -69.8227 |
| 86006430 | -69.8213 |
| 22864 | -69.8089 |
| 12574939 | -69.8077 |
| 8327 | -69.806 |
| 625497 | -69.8045 |
| 7638602 | -69.7624 |
| 7574665 | -69.7557 |
| 71338958 | -69.7311 |
| 76696 | -69.7137 |
| 4537595 | -69.688 |
| 81019 | -69.6734 |
| 12631751 | -69.662 |
| 3853312 | -69.6323 |
| 108771 | -69.5852 |
| 591649 | -69.5657 |
| 235822 | -69.562 |
| 53931115 | -69.5602 |
| 7064878 | -69.5402 |
| 14650 | -69.508 |
| 66504 | -69.5002 |
| 7451902 | -69.423 |
| 30449372 | -69.4165 |
| 131851432 | -69.405 |
| 165832 | -69.3888 |
| 4346802 | -69.3516 |
| 79314 | -69.2939 |
| 2833781 | -69.185 |
| 3316477 | -69.1762 |
| 21423370 | -69.145 |
| 459272 | -69.1312 |
| 66484 | -69.091 |
| 14548391 | -69.0862 |
| 236905 | -69.0517 |
| 77724 | -69.0463 |
| 412443 | -69.03 |
| 8511 | -68.945 |
| 2748264 | -68.9227 |
| 235823 | -68.904 |
| 88943 | -68.8996 |
| 91968 | -68.8287 |
| 4404944 | -68.8148 |
| 522812 | -68.7727 |
| 7164463 | -68.7666 |
| 5484079 | -68.7241 |
| 3127230 | -68.609 |
| 4542191 | -68.5403 |
| 5484533 | -68.4733 |
| 87405 | -68.3799 |
| 2749906 | -68.2263 |
| 10588796 | -68.1403 |
| 175269 | -68.0924 |
| 65731 | -68.0748 |
| 7154852 | -68.0238 |
| 17190 | -68.0172 |
| 8513 | -67.9867 |
| 10778434 | -67.9067 |
| 16206042 | -67.906 |
| 21413448 | -67.8457 |
| 13227198 | -67.7646 |
| 19933616 | -67.5049 |
| 135492599 | -67.3215 |
| 21413435 | -67.3188 |
| 6710 | -67.0453 |
| 2837525 | -67.0085 |
| 27658 | -66.9771 |
| 4311528 | -66.9662 |
| 46494813 | -66.8727 |
| 27657 | -66.842 |
| 11045491 | -66.8417 |
| 92132573 | -66.8356 |
| 71358242 | -66.7782 |
| 6780 | -66.6151 |
| 15555081 | -66.5171 |
| 2847288 | -66.4966 |
| 23680289 | -66.2854 |
| 275870 | -65.8759 |
| 20456763 | -65.6495 |
| 354144 | -64.75 |
| 67209 | -64.0802 |
| 3688907 | -63.702 |
| 7358958 | -61.9614 |
| 3090909 | -61.123 |

**Supplementary Table 2.**
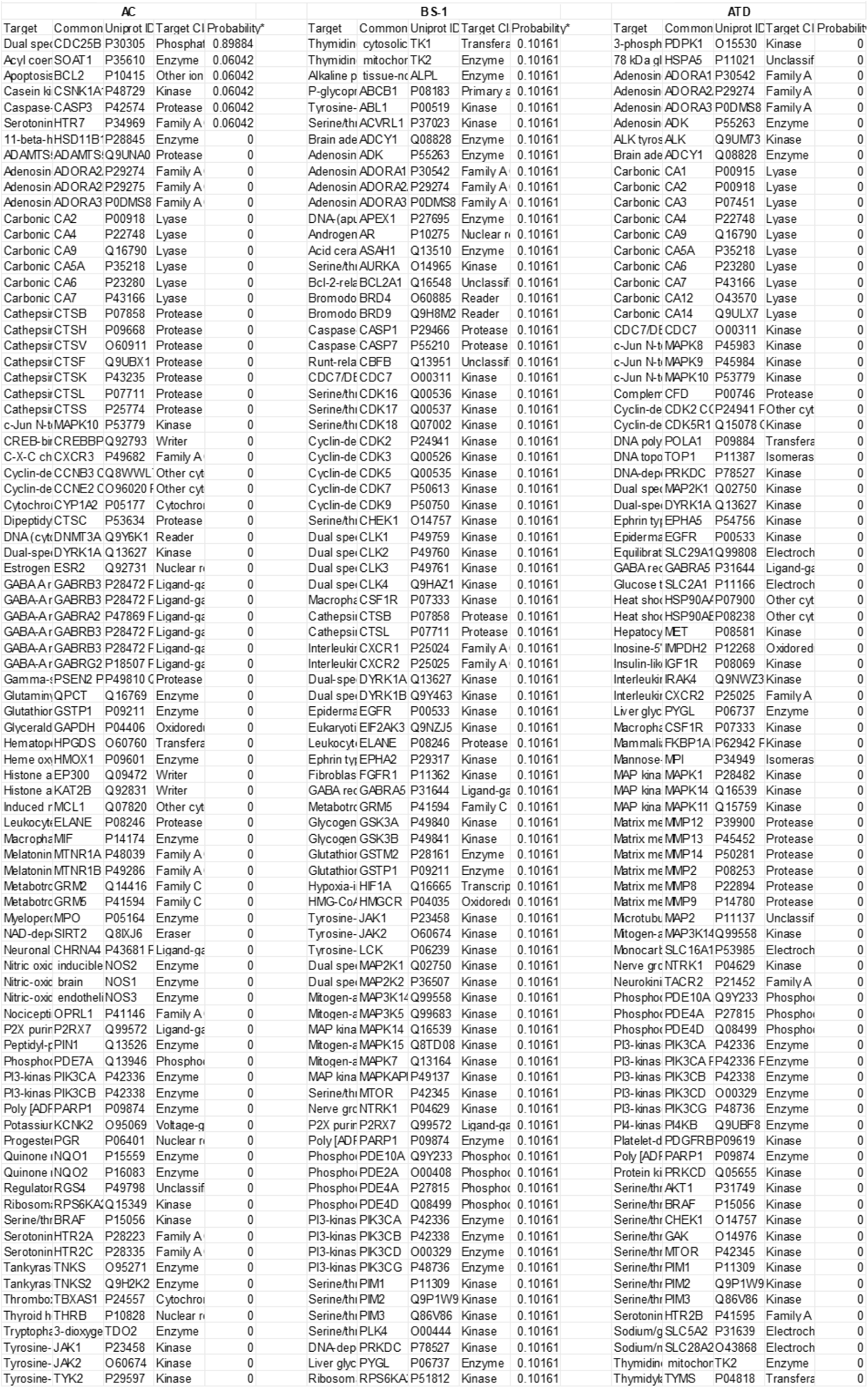
AC, BS-1 and ATD target prediction

